# Learning predictive latent structure from unclustered neural spike trains

**DOI:** 10.64898/2026.11.13.766468

**Authors:** Samuel R. Bray, Kenneth Kay, Christina J. Li, Eric L. Denovellis, Alison E. Comrie, Abhilasha Joshi, Xulu Sun, Alan D. Kaplan, Uri T. Eden, Loren M. Frank

**Author notes:** Correspondence (L.M.F.).

## Abstract

Neural activity patterns enable perception, thought, and action. A powerful approach to understanding these abilities involves identifying low-dimensional, interpretable latent variables that capture the relationship between activity patterns, the information they represent, and the computations they support. Yet current methods often yield variables that are difficult to interpret and require data preprocessing that limits their accuracy and temporal resolution. Here we introduce Contrastive Prediction of Point Process Observations (C3PO), an unsupervised method that learns a predictive, interpretable latent variable representation directly from continuous time neural spiking without requiring spike-sorting or binning. C3PO recovers structure associated with known latent variables across a variety of data sets without access to those variables during training. It identifies dynamics on timescales from tens of milliseconds to hours and reveals previously unreported representations and their dynamics. These results establish C3PO as a powerful tool for uncovering interpretable structure in high-dimensional point-process data sets.

---

The brain enables organisms to perceive their environment, create internal models, make decisions, and execute actions. These abilities depend on coordinated neuron population activity, whose firing patterns reflect information being represented and transformed within a circuit. Often, a relatively small number of generative variables shape population activity^1^. Some are readily interpretable and experimentally accessible, such as delivered stimuli or animal location. Others are not directly observable, instead reflecting internal state, task context, learned value, or other ongoing computations. Moreover, individual neurons can exhibit mixed selectivity to multiple factors^2^, such that the response to any one variable depends on the broader population state and behavioral context. These properties make it more difficult to determine in advance which variables should be controlled, measured, or tested during experiments and analysis, and more difficult to understand the relationship between these variables and neuron activity.

These challenges motivated the development of a broad range of approaches for semi- and un-supervised identification of lower-dimensional structure and dynamics in neural activity. Classical methods such as principal component analysis (PCA) and Gaussian-process factor analysis (GPFA) extract dominant patterns of population covariation, while approaches such as jPCA characterize structured temporal evolution within population states^3,4^. More recent latent-dynamical and representation-learning methods, including LFADS, DFINE, MARBLE, and CEBRA, use more flexible models to capture nonlinear and behaviorally relevant structure in high-dimensional activity^5–8^. These approaches have produced important insights across neural systems but have three important limitations constraining potential discoveries.

First, they depend on spike sorting, where spike events in extracellular recordings are detected and assigned to putative single neurons. This remains a time consuming and imperfect process where many low-confidence spike waveforms are discarded and some fraction are misassigned^9^. Further, accurate real-time spike sorting, particularly for many-electrode probes, is difficult at best, limiting these methods in real-time feedback experiments. By contrast, “clusterless” approaches that avoid spike sorting can enable more accurate decoding^10,11^, and are compatible with real-time feedback^12^.

Second, current methods limit temporal resolution of the inferred features. This constraint arises from binning or smoothing spike trains, restricting conclusions to features slower than the specified bin. Conversely, methods that operate on individual trials typically restrict inference to within-trial timescales^13^, while methods that exploit shared temporal structure across trials may additionally require temporal alignment or warping^14,15^.

Third, these methods require compromises between model flexibility and feature interpretability. Linear approaches provide latent variables that relate directly to neural activity but are often difficult to interpret due to nonlinear tuning, mixed selectivity, and activity that lies on curved manifolds. More flexible nonlinear and deep-learning approaches relax these constraints to capture richer population structure. However, this flexibility often comes at the cost of interpretability: distances and directions in latent space need not correspond straightforwardly to changes in neural activity, and the contribution of individual neurons to a discovered feature can be difficult to determine^16^.

Here we introduce Contrastive Prediction of Point-Process Observations (C3PO), an unsupervised method that addresses these limitations. C3PO identifies interpretable latent variables from unclustered (or clustered) spiking activity, captures time-varying engagement of population activity, and enables direct mapping of latent variables to single neurons—without compromising temporal resolution. We show that C3PO accurately identifies latent features organized by known drivers of neuron tuning including proprioception, position, and internal states, despite having no access to these features during training. We further demonstrate C3PO’s use as a discovery tool by identifying new cross-timescale interactions in prefrontal activity during a complex foraging task. Because C3PO models spikes as point processes rather than pre-binned rates, the extracted features track dynamics ranging from milliseconds to hours. Finally, to illustrate the robustness and scalability of C3PO, we trained and published models for 51 publicly available datasets and show that the inferred contexts replicate findings from the original publications. Together, these results establish C3PO as a powerful and scalable predictive framework for inferring interpretable latent structure from high-dimensional point-process datasets.

## RESULTS

### Model Design

Understanding neural activity involves identifying underlying generative processes. We define *G* as a sufficient statistic of that process such that there exists a non-linear tuning curve, *f*_*j*_(*G*_*t*_), over the generative space predictive of the firing rate of each neuron *j* at time *t*. Neurons fire as stochastic emissions from these conditional rates, recorded as voltage deflections across channels in electrophysiological measurements (Fig. 1A). This results in a sequence of marked, point process observations 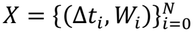 where Δ*t*_*i*_ and *W*_*i*_ are the wait time and waveform feature measurements of the *i*-th observed spike event, respectively, and *N* is the total number of spikes recorded in the experiment. Rather than spike sorting and binning these events into time-averaged firing rates, C3PO directly models this marked point process^17–19^. Inferring *G*_*t*_ from these observations relies on three stages: waveform embedding, context inference, and relative rate prediction (Fig. 1B).

**Figure 1.**
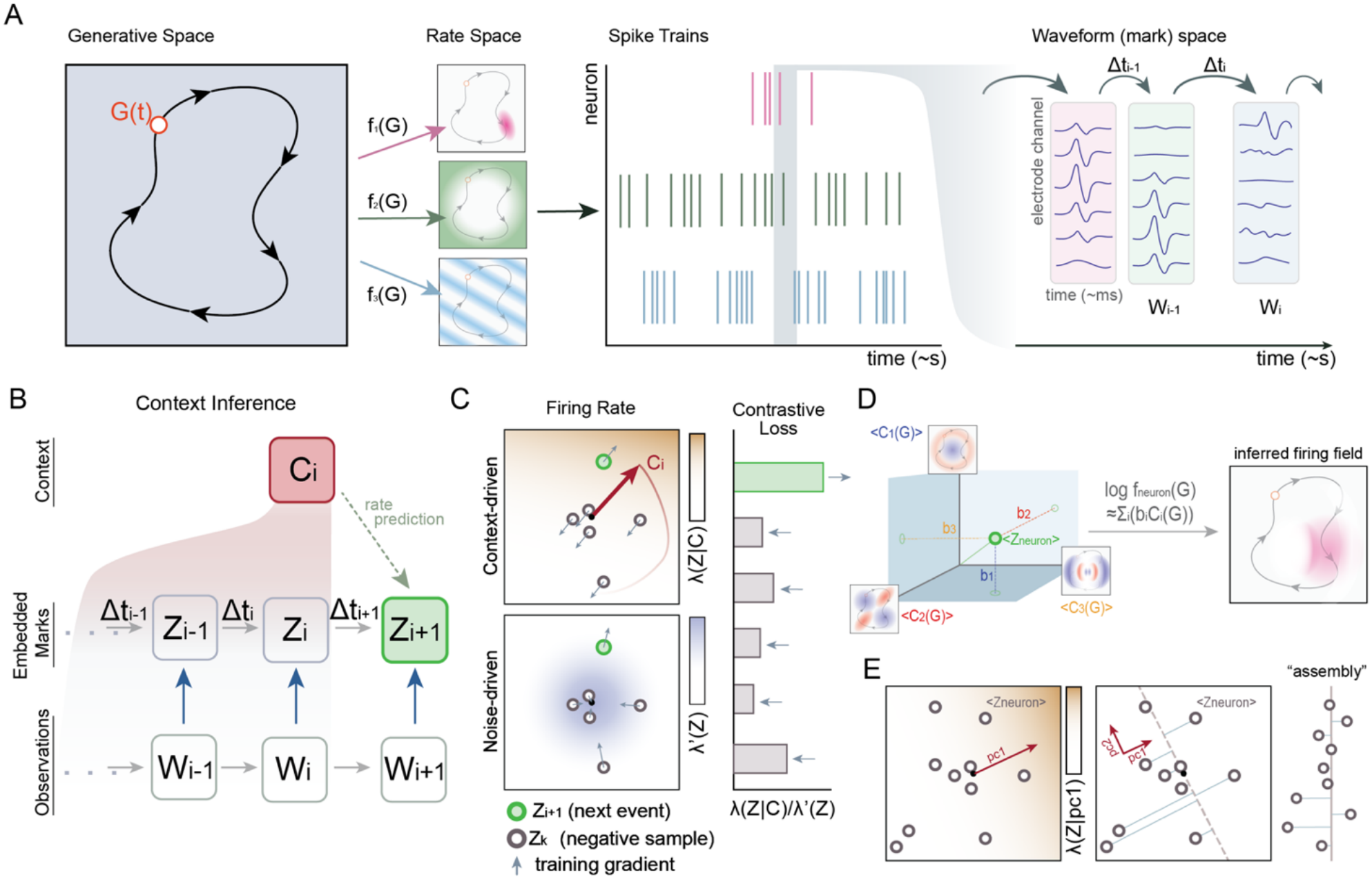
C3PO architecture and interpretation. **A**, Conceptual model of neural observations. A latent generative state G(t) evolves through internal and external dynamics, determining neuronal firing rates through neuron-specific tuning functions f_i_(G(t)). Stochastic spikes produce measured extracellular waveforms separated by inter-event intervals. **B**, C3PO architecture. Waveform-feature vectors (*W*) are independently transformed into mark embeddings (*Z*). A causal neural network integrates preceding embeddings and inter-event intervals to infer a context vector (*C*) that predicts the next event Z_i+1_. **C**, Contrastive training distinguishes the true next event (green) from negative samples drawn from other times in the event sequence (gray). Candidates are scored by the ratio of their context-dependent hazard to a background hazard. The context–mark inner product determines log hazard, whereas the background hazard decreases with embedding magnitude. Arrows indicate gradient updates that promote context alignment with the true event and reduce negative-event scores by shrinking negative embeddings toward the origin. **D**, Context components form nonlinear functions of the generative state (insets). A neuron’s mean waveform embedding supplies loadings on these functions, whose weighted sum predicts its log firing-rate tuning. **E**, Projections of neuronal embeddings onto a context direction (here the first principal component of C) define signed weights for a co-modulated neural assembly.

During embedding, each waveform *W*_*i*_ is projected from its high dimensional measurement (∼10^2^ – 10^4^ for high-density probes) to a lower dimensional mark *Z*_*i*_ through a neural network (Fig. S1). Here we use the peak amplitude of the signal on every electrode during the spike as input features, but any informative features (including sorted neuron id’s) can be used. After embedding, the observations form a new series 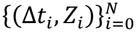 .

Next, we infer the state of the generative system from the history of embedded events using a causal framework. The key is including sufficient information about past spiking to inform current state. We use a dilated convolutional network (Wavenet^20^) that takes information from the previous (∼10^3^) spikes to estimate the current context, although any network that can include such history can be applied (e.g., recurrent neural networks or transformers). This results in a “context” variable *C* that is optimized to capture information about the generative state from prior spiking activity.

To ensure an interpretable relationship between *C* and spiking, we define a model structure where the log firing rate is linearly determined by how aligned the context vector, *C* is with the mark, *Z*, for a spike. Specifically, we model the conditional firing process with a joint mark intensity function *λ*(*t, Z*_*j*_|*C*_*i*+_) that defines the instantaneous rate of events based on their embedded waveform *Z*_*j*_ during the interval (*t*_*i*−1_, *t*_*i*_]. We model events as a marked Poisson process, though any parameterizable point process can be used, and set:

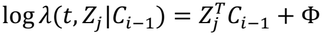

where Φ is a leading coefficient that negates in the loss function (see Methods). This allows us to interpret the context as a vector in mark space pointing toward neurons with high conditional firing rates.

Training simultaneously optimizes the mark-embedding and context networks. As a standard likelihood-based approach is not tractable (see Methods), we introduce an approach derived from Contrastive Predictive Coding (CPC) and Noise Contrastive Estimation (NCE)^21,22^, and adapted for point process data. Intuitively, the model is tasked to identify the true event (the next waveform *Z*_*i*_) from a set of negative events sampled from elsewhere within the data 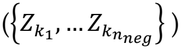 based on the ratio of the waveform’s firing rate at this time (i.e., the conditional firing rate *λ*(*t, Z*_*j*_|*C*)) relative to the “noisy” unconditioned firing rate of that waveform, *λ*′(*Z*_*j*_). (Fig. 1C; see Methods).

The noisy firing rate term describes spiking independent of (and unpredictable by) the system state. We model this activity as a Poisson process with a log rate inversely scaled by the embedded waveform magnitude:

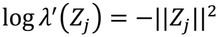

This regularizes the embedded space—events will maintain magnitude if they can be reliably distinguished by a particular context, and events that are not predicted by the context will collapse to the origin (Fig. 1C).

Gradient updates align the context and the true event (to maximize conditional likelihood), while pushing negative samples away from the current context (Fig. 1C). Critically, this forces the model to learn contexts that differentiate times when different subsets of neurons fire. Simultaneously, the embedded waveform space clusters co-occurring spikes together, such that a single value of the context predicts the population activity.

### C3PO Interpretation

#### Context as a basis set over generative space

This design creates key relationships for model interpretability. The inferred context *C* summarizes the system state predictive of firing rates—approximating the information carried by *G* in our conceptual model (Fig. 1A). To do so, the context components *C*^(*j*)^ must systematically vary with *G*, giving an average context response ⟨ *C*^(*j*)^(*G*)⟩ (Fig. 1D).

At the same time, *C* directly predicts neuron firing rates *f*_1_(*G*) such that:

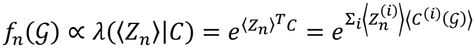

where ⟨*Z*_1_⟩ is the average embedding of all spikes from this neuron. This means the components of *C* describe a non-linear basis set over the generative space (Fig. 1C). In turn, the components of a neuron’s embedded waveform ⟨*Z*_1_⟩ comprise the appropriate loadings on these bases to approximate a neuron’s firing over time (Fig. 1D).

#### Directions in context space describe assemblies of neural activity

Complementarily, directions in context space describe patterns of population activity. Context dynamics along a direction 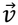predict firing rate modulations scaled by the magnitude of neuron embeddings along that axis:

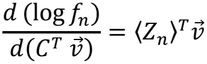

Weights of projected neuron embeddings describe participation in firing rate co-modulation, e.g., a neural assembly (Fig. 1E). Different directions in context space allow C3PO to describe activity of neurons participating in multiple modulations with mixed selectivity.

### Toy model of periodic neural population activity

For initial validation we constructed a synthetic dataset of neural activity driven by a single periodic function *G*_*t*_ = *θ*(*t*) with a period of 3 seconds (Fig. 2A). We define 48 neurons with nonlinear tuning curves,*f*_1_(*θ*), over this latent phase, each with a preferred phase. Additionally, we include 16 non-informative noise units, with flat tuning curves independent of *θ*. To mimic extracellular recordings, each neuron is assigned a template waveform across 128 recording channels. Individual events are constructed by adding noise to these templates.

**Figure 2.**
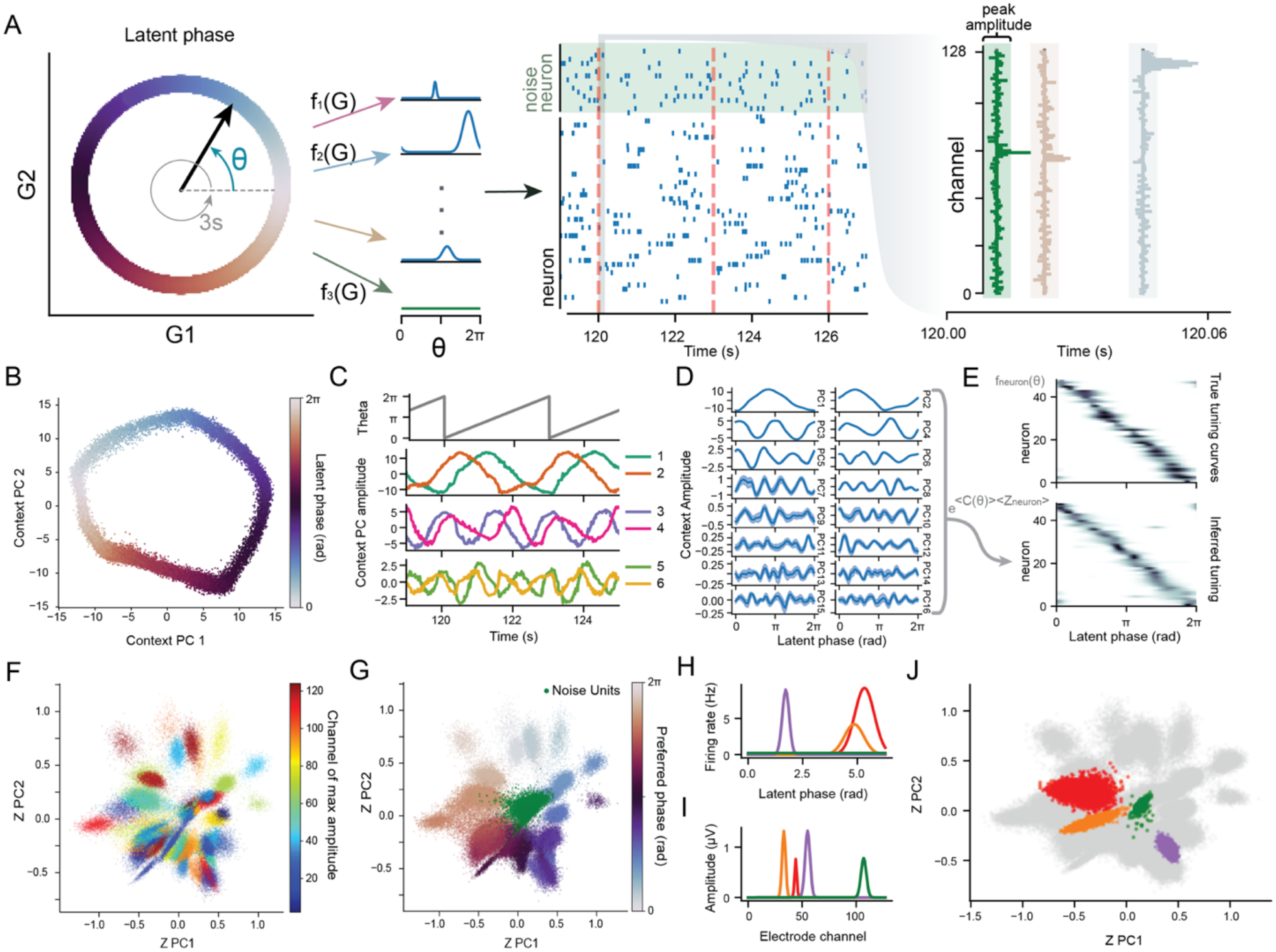
C3PO recovers latent dynamics and neuronal tuning from unclustered spike events. **A**, Synthetic marked spike trains are generated over 5,000 s from 48 phase-tuned neurons and 16 untuned noise neurons (green). Panels follow the organization in **Fig. 1A**. Left, latent phase advances with a 3-s period and determines neuronal tuning; middle, example spike raster; right, noisy 128-channel waveform-feature vectors associated with individual events. A 16-dimensional C3PO model was trained without access to latent phase or neuron identity. **B**, Context projected onto its first two principal components (PCs), colored by latent phase. **C**, Example time courses of actual latent phase (top) and the first six context PCs. **D**, Median and interquartile context-PC values as a function of latent phase. **E**, Ground-truth (top) and inferred (bottom) tuning curves for the 48 phase-tuned neurons, ordered by preferred phase. Inferred curves were calculated by exponentiating the inner product of each neuron’s mean mark embedding and the phase-conditioned mean context, then normalizing by their mean across phase bins. **F**,**G**, Mark embeddings projected onto the first two context principal axes, colored by the channel of peak waveform amplitude (**F**) or the emitting neuron’s preferred phase (**G**). Green points in **G** denote untuned neurons. **H-J**, Tuning curves (**H**), waveform-feature profiles across recording channels (**I**), and mark embeddings (**J**) for four example neurons. Colors identify the same neurons across panels; other events are gray in **J**.

#### Inferred context constructs a basis set over generative phase

We trained a 16-dimensional C3PO model on these data. The number of dimensions was chosen as sufficient to capture the key features of the data (Fig S2A) and higher dimensional models yielded similar results.

Context values were transformed using principal components analysis (PCA). Importantly, the linear rate prediction within C3PO permits a linear change of basis without altering context meaning. PCA features thereby naturally capture directions of shared neural firing rate variability.

To examine the learned geometry, we plotted the first two PCs of *C* at each timepoint, colored by *θ* (Fig. 2B). This gave a clear ring manifold, organized by generative phase. Notably, higher PCs also display periodic activity, with increased frequency (Fig. 2C). We confirmed this by calculating the average context values across *θ*. Strikingly, this produced pairs of phase-shifted context components with an approximate frequency increase of 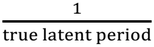 between pairs (Fig. 2DI, S2 B-C). These effectively approximate a Fourier-basis over *θ*, and thus a linear weighting of the PCs can approximate arbitrary periodic firing rate functions for each neuron (Fig. 1D).

To verify, we calculated each neuron’s average embedded waveform ⟨*Z*_*neuron*_⟩ and used them to infer tuning curves log *f*_*n*_(*θ*) ∼⟨*Z*_*n*_⟩⟨*C*(*θ*)⟩^*T*^. These tuning curves, inferred only from unclustered firing data, show remarkable alignment with the true values (Fig. 2E).

#### Learned waveform embeddings organize according to tuning information

The waveform embedding space (*Z*) acts as a bottleneck layer retaining information on tuning patterns while discarding measurement details. To demonstrate this feature, we compared spike embeddings with their location on the measurement probe (Fig. 2F). While events from the same neuron do cluster, there is no global organization on waveform feature. By contrast, when we compare embedded waveforms to the preferred firing phase of the emitting neuron, we see a clear global organization based on tuning (Fig. 2G). This agrees with interpretation of neural assemblies, where C3PO organizes waveform embeddings based on co-modulation (Fig. 1E). Because “noisy” neurons lack modulation by the context, embedded waveforms from these units cluster near the origin (Fig. 2G). To further illustrate this, we compared tuning curves, waveforms, and embeddings of sample neurons (Fig. 2H-J). Again, we see that embedding proximity is based on tuning information rather than waveform similarity.

### Latent variable identification from Macaque S1

As an initial validation in neural recordings, we applied C3PO to sorted spikes from primary somatosensory cortex (S1, area 2) in a rhesus macaque during active reaches and passive arm perturbations^23^ (Fig. S3A,B). Without behavioral annotations during training, C3PO recovered context trajectories organized by movement direction and whether movement was self-driven (Fig. S3C). Supervised decoders fitted to the learned context reconstructed hand trajectories and classified eight reach directions with 84% accuracy on held-out trials (Fig. S3D,E). This level of accuracy is comparable to that previously reported using a supervised version of CEBRA^8^. Hand, joint, and muscle-derived variables were also decodable (Fig S3F). In particular, there was high decodability for the biomechanically related x-directed hand movement and lengths of the pectoral muscles that was reflected in the learned geometry (Fig. S3F). These findings are consistent with the whole-arm proprioceptive organization reported in the original study^23^, demonstrating that C3PO recovers interpretable movement-related structure without specifying behavioral variables during representation learning.

### Latent variable discovery from rat entorhinal cortex

Having established that C3PO can recover behavioral structure from neural data, we asked whether it could resolve multiple, simultaneously expressed modulators of population activity. To do so, we trained a 16-dimensional C3PO model on a publicly available Neuropixels recording of 460 sorted neurons from the rat medial entorhinal cortex (MEC) and adjacent parasubiculum during open field foraging^24^ (Fig. 3A). These regions provide a useful test because neurons are heterogeneously driven by several well-characterized variables, including an ∼8Hz theta rhythm, head direction, and spatial position, resulting in mixed selectivity at the neuron level (Fig. 3B)^25–27^.

**Figure 3.**
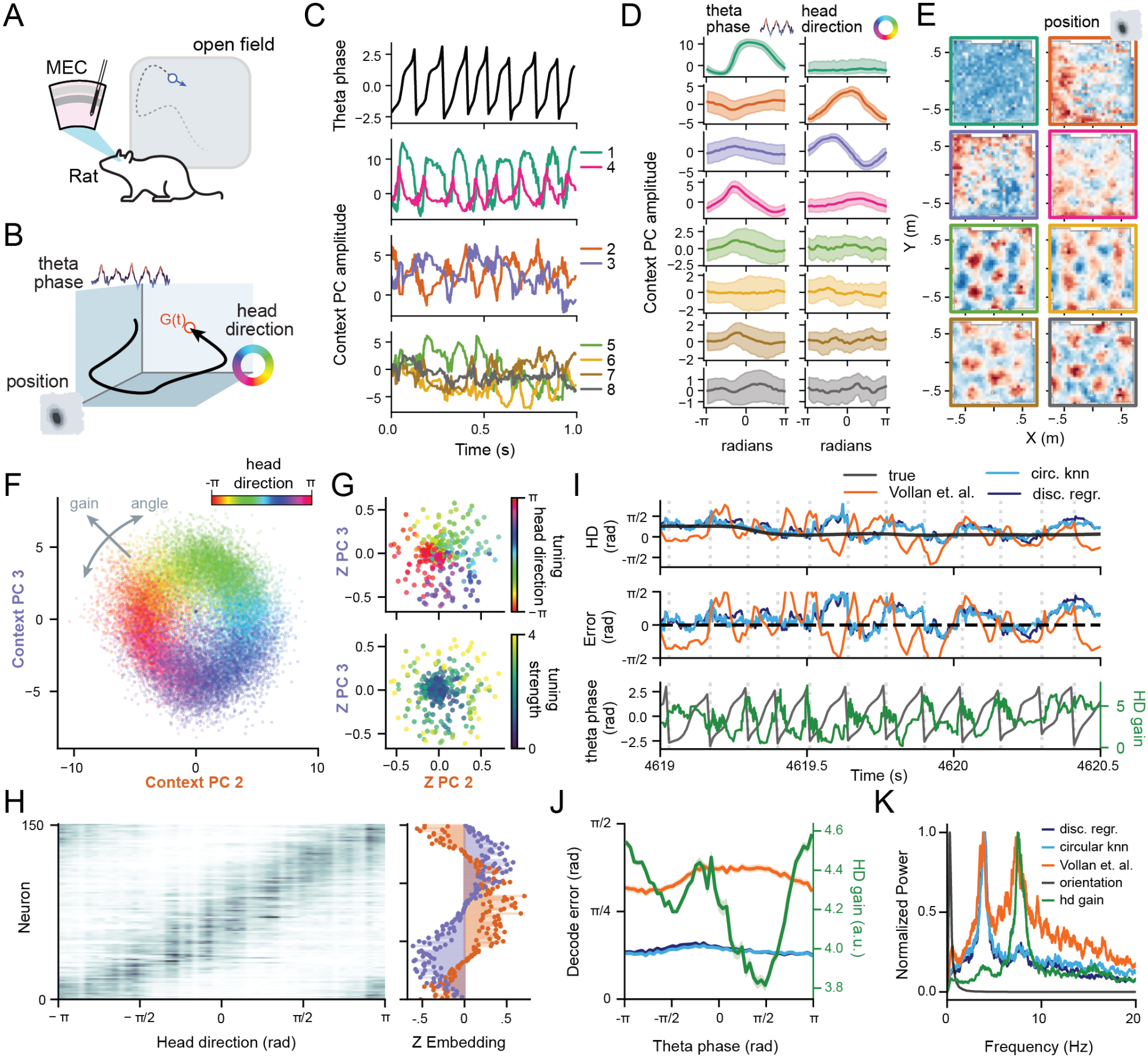
C3PO isolates multiple representations in entorhinal population activity. **A**, Neuropixels recording from medial entorhinal cortex (MEC) and adjacent parasubiculum during open-field foraging, comprising 460 sorted neurons from one rat (Vollan et al., 2025). **B**, A cartoon of the generative space for MEC (G(t)) highlighting known drivers of activity, including theta phase, head direction, and spatial position. **C**, Example trace of theta phase (top) and the first eight PCs of C3PO context. **D**, Context-PC values conditioned on theta phase (left) or head direction (right). Lines and shading indicate median and interquartile values, respectively. **E**, Spatial maps of median context values for PCs 1–8, arranged in row-major order; border colors match **C, D**. Each map uses a symmetric color scale spanning its maximum absolute value, with red and blue indicating negative and positive values, respectively. **F**, Context projected onto PCs 2 and 3, colored by measured head direction. Context angle and gain denote the polar angle and radial magnitude within this plane. **G**, Learned neuronal embeddings projected onto the same PC axes, colored by preferred head direction (top) or tuning strength (bottom). Preferred direction is the angular bin with maximal firing rate; tuning strength is the base-2 logarithm of the peak-to-median firing-rate ratio across angular bins. **H**, Head-direction tuning curves normalized by each neuron’s mean across angular bins (left), alongside neuronal embedding weights on PC2 (orange) and PC3 (purple; right). Neurons with embedding magnitude greater than 0.2 in this plane are included and ordered by embedding angle. **I**, Example head-direction dynamics. Top, measured head direction (gray), five-fold cross-validated C3PO predictions from multinomial logistic regression (dark blue) and circular nearest-neighbor regression (light blue), and the internal-direction estimate supplied by Vollan et al. (orange). Middle, signed circular decoding errors relative to measured head direction. Bottom, theta phase (gray) and context gain (green). Vertical dashed lines mark theta-phase cycles. **J**, Median absolute decoding error (left axis) and median context gain (right axis) as functions of theta phase; colors follow **I**. **K**, Frequency-weighted power spectra of signed decoding error, measured head orientation (gray), and context gain (green), each normalized to its maximum. Decoding errors exhibit a prominent approximately 4-Hz component, whereas gain is predominantly modulated near the 8-Hz theta frequency.

The inferred context captures these variables, identifying structured fluctuations spanning sub-second (theta) and behavioral timescales (Fig. 3C). Context PCs 1 and 4 are phase-shifted basis functions over theta phase, indicating C3PO identified theta-timing differences across the neural population (see Sup. Fig. 4 for additional details on theta phase modulation). By contrast, PCs 2 and 3 are a separate pair of phase-shifted functions spanning head direction (Fig. 3D). These pairs provide low-dimensional representations of these circular variables. Higher-order PCs captured spatial position, where PCs 5-8 exhibited hexagonal grid-like response patterns across the open field, with different spatial offsets across components (Fig. 3E).

**Figure 4.**
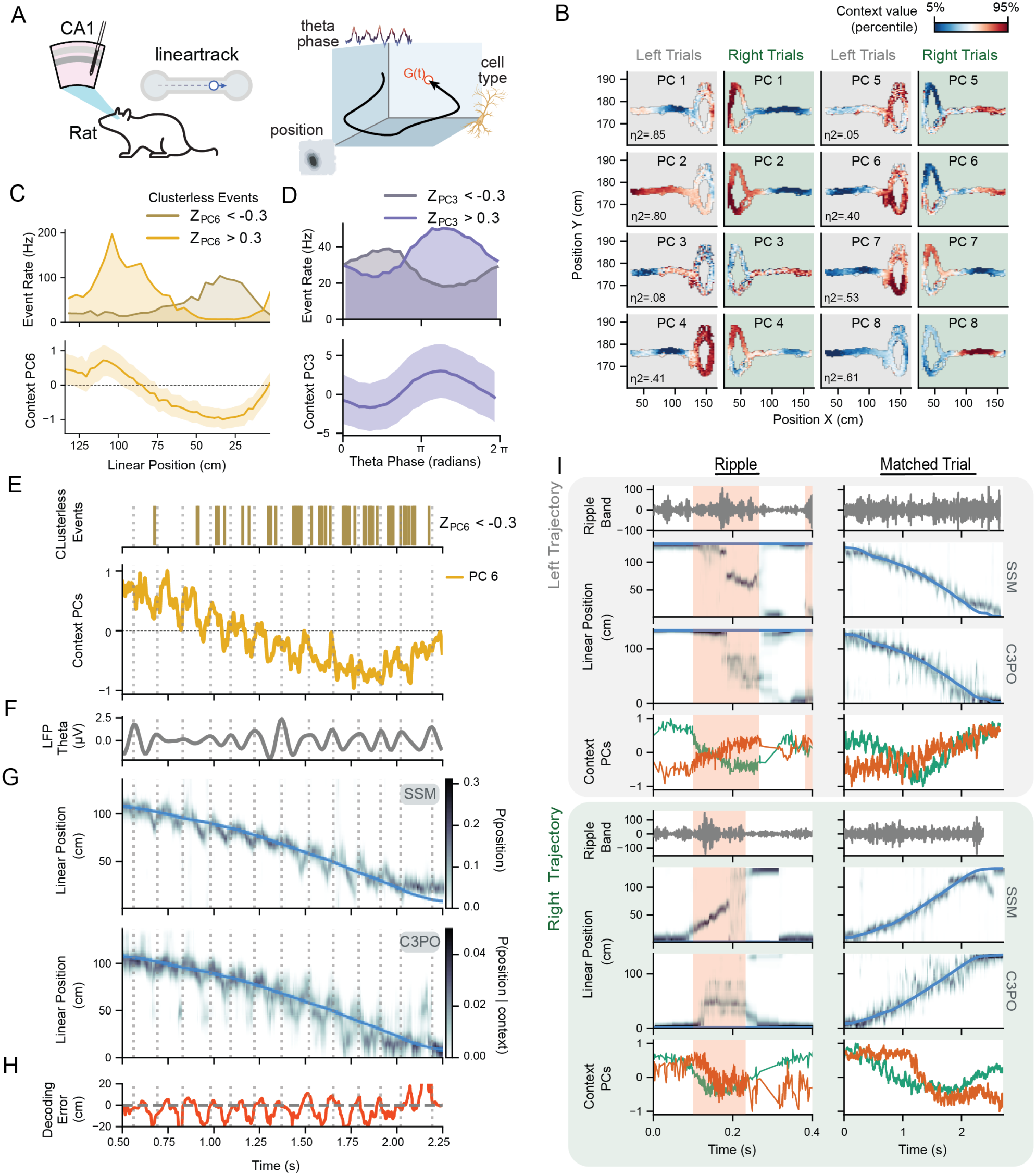
C3PO captures repeated population structure across timescales in hippocampal area CA1. **A**, Schematic of tetrode recordings from dorsal CA1 in a rat traversing a 125-cm linear track, illustrating position, theta phase and cell type as sources of population structure^28^. **B**, Mean values of the first eight context principal components (PCs) across position, shown separately for leftward and rightward trials. Color limits span the 5th–95th percentiles of spatial-bin means for each PC. Annotations report squared correlation ratios for the joint dependence on linear position and travel direction. **C**, Spatial correspondence between context and waveform embeddings during leftward running. Top, event rates for waveform embeddings with projections onto context PC6 below ™0.3 (olive) or above 0.3 (gold). Bottom, median, and interquartile values for context PC6 versus position. High magnitude context corresponds to locations with elevated rates of clusterless events with the same embedded direction. **D**, Theta-phase dependence of events with PC3 projections below ™0.3 (gray-purple) or above 0.3 (purple; top), and median context PC3 with interquartile shading (bottom). Opposite context excursions correspond to the preferred theta phases of the respective event groups. **E**, Example leftward traversal showing events with PC6 projections below ™0.3 (top) and the corresponding context PC6 trace (bottom). **F**, Theta-band LFP (5–11 Hz) during the traversal in E. Vertical dotted lines across E-H mark theta-LFP maxima. **G**, Position probabilities from the clusterless state-space decoder of Denovellis et al. (top) and a multinomial logistic decoder of C3PO context evaluated using fivefold temporal cross-validation (bottom). Blue curves indicate measured position. Both representations show theta-associated spatial sweeps. **H**, Signed C3PO decoding error, calculated as the posterior-weighted mean position minus measured position. **I**, Representative leftward (upper group) and rightward (lower group) ripple trajectories paired with traversals in the corresponding direction. Within each group, rows show ripple-band LFP, state-space position probabilities, C3PO position probabilities and context PC1 (green) and PC2 (orange). Pink shading marks detected ripple intervals; blue curves show measured position. Context traces are centered and normalized independently within each window and smoothed over 11 spike events. Similar context transitions unfold over substantially shorter intervals during ripples, consistent with temporal compression of spatial population dynamics.

#### Deriving single neuron head-direction tuning from C3PO context

C3PO also provided readily interpretable representations of head direction. Plotting the two head-direction-sensitive context components (PCs 2 and 3) revealed a ring-like structure organized by head direction (Fig. 3F). We defined the angular position within this plane as the head-direction context angle, and its radial magnitude as the head-direction context gain (Fig. 3F). The angle reflects which directional population pattern is expressed, whereas the gain quantifies the strength of pattern activation.

Because context and mark embeddings share a predictive space, neurons tuned to a particular head direction should have embeddings aligned with the same direction-associated context. Similarly, neurons with stronger head-direction modulation should have larger projections within this plane. We therefore compared each neuron’s learned embedding on the PC2–PC3 axes with its head-direction tuning calculated from raw spikes.

Neuron embedding angles closely matched the direction at which each neuron fired maximally (Fig. 3G, top). Moreover, neurons with stronger head-direction tuning lay farther from the origin, whereas weakly tuned neurons clustered near the center (Fig. 3G, bottom; Fig. S4A, R^2^=0.71). This captures population-scale organization: ordering neurons by their embedding angle corresponds to an orderly progression of their head direction tuning curves (Fig. 3H). Thus, C3PO captured both population head direction representations and individual neuron participation in this feature.

#### Rapid, alternating head direction dynamics from C3PO context

We next asked whether context dynamics reflected fast directional fluctuations reported in the original study, which showed that the MEC population did not purely track the measured head direction. Instead, the decoded population direction often shifted to opposite sides of the head axis on successive theta cycles, aligning with left–right spatial sweeps expressed by grid-cell populations^24^.

We also found this structure in C3PO’s context by constructing cross-validated decoders of head direction from context. On longer timescales, the decode tracked the animal’s actual orientation (Fig. 3I, top). On theta timescales, we observed alternating decoding errors across successive theta cycles (Fig. 3I, middle) similar to the original paper (Fig. 3I, orange), which exhibited a prominent 4-Hz component corresponding to a pattern repeating every two theta cycles (Fig. 3K).

C3PO’s geometry provided additional insight into these fluctuations. The head-direction context gain varied strongly across individual theta cycles (Fig. 3I, bottom) and was maximal at phases when decoding error was smallest (Fig. 3J). This suggests current head direction is encoded most strongly at a specific theta phase, with the subsequent shift toward non-actual directions engaging weaker coding. Correspondingly, head-direction gain showed prominent theta-frequency modulation, whereas directional error contained a 4-Hz component from alternating left–right deviations (Fig. 3K).

#### Comparison to existing methods

To compare these results with other latent-variable approaches, we applied CEBRA and LFADS to the same MEC recording and repeated the PCA and head-direction decoding analyses above. Both methods recovered theta, head-direction, and spatial modulation, though with more mixing of features than the C3PO embedding (Figs. S5 and S6, A-D). Both methods also produced latent variables capable of decoding head direction (Figs. S5 and S6, E-F) and associated theta-timescale errors, though only the LFADS embedding enabled decoding performance comparable to C3PO (Fig. S7). Together, these comparisons demonstrate C3PO achieves state-of-art performance in extracting behaviorally informative representations and rapid directional fluctuations, while also identifying direct relationships between population features and individual-neuron tuning.

### Clusterless latent variable discovery from hippocampus

Unlike alternative methods, C3PO also enables discovery of population structure without spike sorting. To illustrate “clusterless” feature discovery, we analyzed previously published rat recordings from 30 tetrodes targeting dorsal CA1 during linear track^28^ (Fig. 4A). We represented each threshold-crossing event by its peak amplitude on each channel and trained a 20-dimensional C3PO model on these marked point-process observations (see Methods). Here our goal is to identify known modulators of CA1 activity, including position and theta rhythm during locomotion^29,30^, and their interactions. Specifically, as an animal moves through a pyramidal cell’s place field, its spikes occur at progressively earlier phases of theta—a phenomenon known as phase precession^31,32^. At the population level, this results in characteristic “theta sweeps” where the position represented by neural activity quickly progresses through locations behind and ahead of the animal within single theta cycles^33–35^.

#### C3PO identifies known generative variables from clusterless marks

We calculated the average context value as a function of position and direction of travel. Many components showed strong spatial localization, with distinct regions of positive and negative activity during running intervals (Fig. 4B). Changes in position-sensitive context PCs corresponded to different rates of waveform events embedded on these axes (Fig. 4C), indicating C3PO identified spatially driven modulations for both individual waveforms and population activity.

C3PO context also showed prominent theta modulation, with power spectra revealing peaks near 8Hz in nearly all PCs, as well as power on behavioral timescales (Fig. S8A). PC3 in particular was more selectively dominated by theta-frequency activity. Averaging over theta phase revealed an approximately sinusoidal basis across the cycle (Fig. 4D), indicating that C3PO had captured theta modulation without LFP information during training. This organization is consistent with differing timing of different CA1 cells within theta oscillations^29,36^. Notably, known cell-type driven firing patterns were identified without clustering or unit labeling.

#### C3PO position context precesses relative to theta phase

Several PCs also showed interactions between position and theta phase (Fig. 4E, F, Fig. S8A). To illustrate this, we trained a linear position decoder from the PCs (Methods). For reference, we compared decoding results with those from an established supervised clusterless state-space decoder applied to the same waveform events^37^. Both approaches successfully tracked position over behavioral timescales (Fig. 4G).

At finer timescales, the posterior decoded from C3PO context repeatedly departed and returned to the measured position within individual theta cycles (Fig. 4G, H). Individual theta cycles show sequential modulation of context components, explaining the linear-decodability of this sweep trajectory (Fig. S8B). The decode error showed a power-spectrum peak near 8 Hz (Fig. S8C) and systematic variation across theta phase (Fig. S8D), confirming that the C3PO extracted the interaction between position and theta generative features. Individual context variables also exhibited structured progression in joint theta-position space during movement (Fig. S8E), corresponding to shifting phase preference of waveforms with loadings along that axis (Fig. S8F). These results show that C3PO can infer multi-timescale features of single cells and population activity without any spike sorting labels.

#### C3PO sequence compression in replay events

Finally, we tested if C3PO representation could recapitulate spatial sequences expressed on still faster timescales. During pauses in movement and sleep, hippocampal populations generate brief sharp-wave ripple (SWR) events in which place-cell sequences associated with spatial experience can be re-expressed in a temporally compressed form, termed replay^38,39^. Spatial population patterns expressed over seconds during locomotion can unfold within tens of milliseconds during immobility, providing a stringent test of the point-process formulation of C3PO.

We applied the same linear decoder to C3PO context and compared it to the state space model. During individual SWRs, the reference decoder exhibited rapid replay trajectories across the track, and the C3PO probe recovered corresponding changes in represented position (Fig. 4I, left), although the C3PO representations were less spatially concentrated, likely as a result of the lack of an explicit continuity model. Nevertheless, these decoded trajectories were accompanied by rapid, ordered transitions in the C3PO context itself (Fig. 4I). Comparing replay events with direction-matched behavioral traversals (Fig. 4I, right) showed that spatial context components recruited over seconds during locomotion were re-expressed in a similar pattern during the ripple event.

### Latent variable discovery from rat prefrontal cortex

Among brain regions, prefrontal cortex shows exceptionally heterogeneous neural responses, making interpretation of its functions and operations an ongoing challenge. Previous studies have identified neural activity organized by representations of behavioral sequence progression^40–42^ and history-dependent signals associated with foraging decisions^13,43,44^. On longer timescales, prefrontal activity can change with task utility as animals approach task disengagement^45^. To test C3PO’s ability to extract features from this rich population activity, we analyzed high-density rat medial prefrontal cortex (mPFC) recordings during a previously described depletion–repletion foraging task where the relative value of staying or switching depended on recent experience^13^ (Fig. 5A,B).

**Figure 5.**
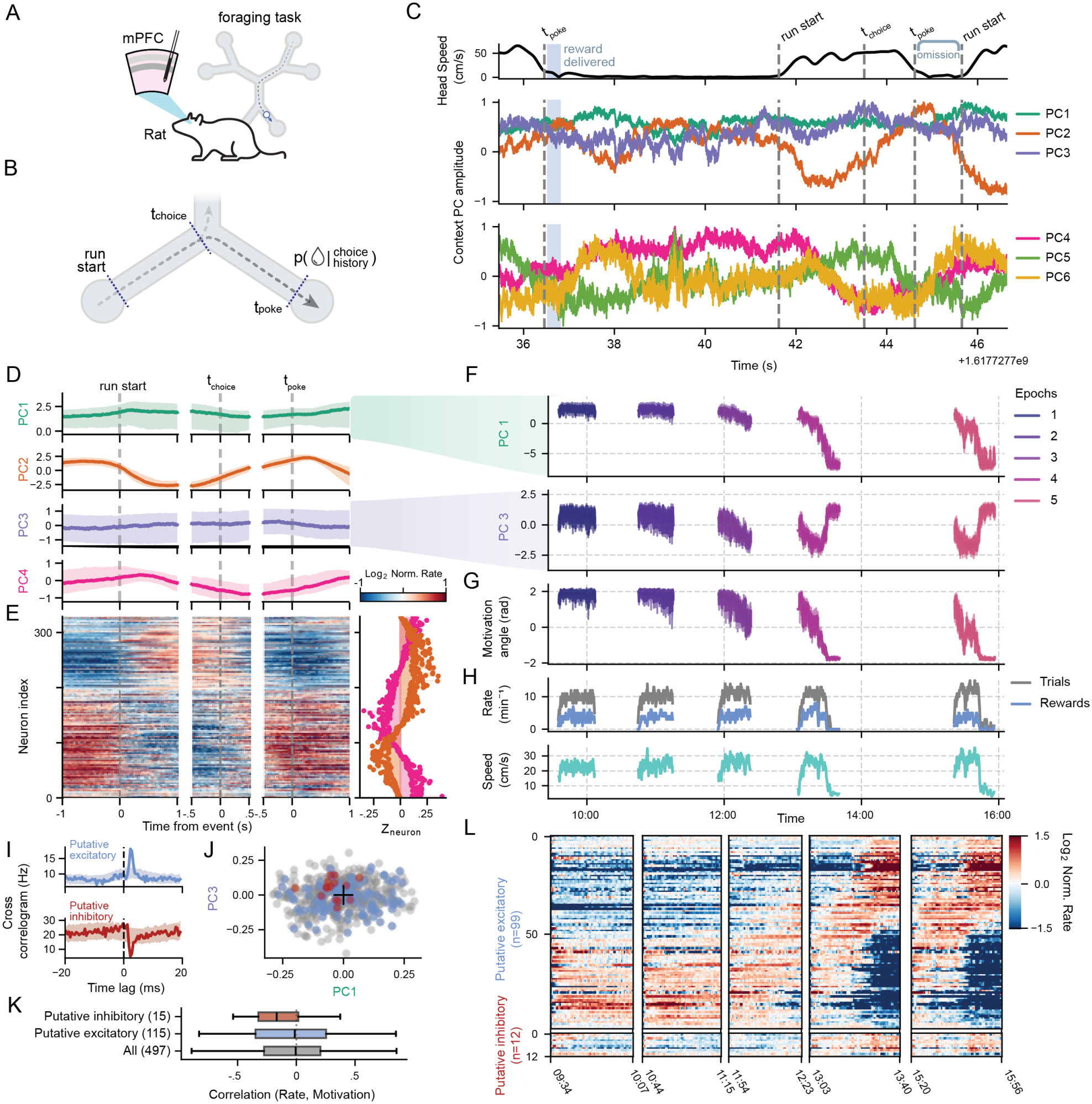
C3PO identifies trial progression and a slow prefrontal population state associated with task engagement. **A**, Medial prefrontal cortex (mPFC) recordings using high-density polymer probes in a rat performing a depletion–repletion foraging task with three patches, each containing two reward ports^13^. **B**, Trial reference events: departure from the preceding port (run start), passage through the choice point (*t*_*choice*_), and entry into the destination port (*t*_*poke*_). Reward probability depended on recent patch choices. **C**, Example head-speed trace and the first six context principal components (PCs). Dashed lines indicate reference events; blue shading marks reward delivery. **D**, Median and interquartile ranges of event-aligned context values across non-switch trials; shading indicates interquartile ranges. PCs 2 and 4 show phase-shifted modulation across successive trial stages, whereas PCs 1 and 3 show comparatively little event-aligned modulation. **E**, Left, PC2 and PC4 loadings of 329 sorted-neuron embeddings with projection magnitude above 0.1 on these axes, ordered by embedding angle. Right, corresponding trial-averaged firing rates, smoothed with a 15-ms Gaussian kernel and displayed as log_2_ ratios to each neuron’s mean rate during aligned windows. Embedding order reveals sequential firing across trial stages. **F**, Context PC1 and PC3 across five task epochs, colored by epoch. Curves show medians within 3 s windows; shading indicates interquartile ranges. Large excursions emerge late in epochs 4 and 5, with values returning toward their earlier range after intervening rest. **G**, Angular position of context in the PC1–PC3 plane, used as an operational neural index of “motivation.” **H**, Trial rate (gray), reward rate (blue), and mean head speed (orange) summarized over 60-s windows. Neural state transitions begin before the cessation of task performance in the later epochs. **I**, Example spike cross-correlograms showing short-latency peaks (blue) and troughs (red), used to infer putative excitatory and inhibitory interactions using spike-time jitter controls adapted from Fujisawa et al. (2008). Shading indicates the pointwise range of jittered cross-correlograms. **J**, Sorted-neuron embeddings projected onto the PC1–PC3 plane. Putative excitatory neurons are blue, putative inhibitory neurons red, and remaining neurons gray; the black cross marks the origin. **K**, Distributions of trial-by-trial Pearson correlations between neuronal firing rate and the neural motivation index across neurons. Boxes indicate interquartile ranges, center lines medians and whiskers the most extreme values within 1.5 interquartile ranges of the box. **L**, Relative firing rates across all five epochs for 99 putative excitatory and 12 putative inhibitory neurons with PC1–PC3 projection magnitude above 0.1. Neurons are ordered within each class by embedding angle, with row order preserved across epochs. Rates are calculated in 60-s windows advanced in 10-s steps and expressed as log_2_ ratios to each neuron’s mean across epochs. The activity progression accompanies the slow context transition, with putative inhibitory neurons tending toward higher firing at lower motivation-index values.

To leverage both the sorted and raw electrophysiological data available for this dataset, we developed an approach to retain dense observations from clusterless events while preserving an explicit connection to neuron identities and annotations. To do so, we implemented a hybrid C3PO encoder mapping sorted and clusterless events into a common embedded space (Methods; Fig. S1).

#### Task Progression Bases in prefrontal activity

We trained the hybrid model across five run epochs and examined the inferred context in relation to trial structure (Fig. 5C, D). We noted that PCs 1 and 3 did not show trial-based modulation, a point we return to below. By contrast, PCs 2 and 4 showed structured fluctuations around run initiation, choice, and reward port entry, forming approximately sinusoidal basis functions over path progression. (Fig. 5B, D).

With the hybrid model, we then projected sorted neurons’ mark embeddings onto the task-progression PC’s, selected neurons with strong projections, and ordered them by embedding angle, which predicts an ordered progression of preferred activity across trial stages (Fig. 5E, right). We validated this by plotting the relative firing rates around the same behavioral events, retaining embedding-defined ordering. This revealed sequential activation across trial stages (Fig. 5E, left).

#### C3PO identifies slow internal state preceding task disengagement

PCs 1 and 3 showed little modulation across trial stages (Fig. 5D) or individual trials (Fig. 5C), suggesting a structure on longer timescales (Fig. 5F). Dynamics in the first three epochs showed relatively modest shifts, with substantially larger excursions at the end of epochs 4 and 5. Context values reset toward their earlier range at subsequent run epochs, which were separated by rest sessions. We trained equivalent sorted-only and clusterless-only C3PO models and confirmed these dynamics were not an artifact of the mark feature representation and could be identified by C3PO without spike sorting (Fig. S9A-C). This longer timescale structure was most readily identifiable in models including the unclustered spikes (Fig. S9D)

We quantified trial rate, reward rate, and velocity through run epochs (Fig. 5H). During epochs 4 and 5, the animal disengaged from the task prior to the experimentally imposed 300-trial limit, despite receiving rewards at a similar rate to previous epochs. These behavioral changes were preceded by the large excursions in the PC1–PC3 context plane. Notably, PCs 1 and 3 showed little systematic variation between running and reward-port occupancy within individual trials (Fig. 5C, D), suggesting they describe a slow population state associated with changing task engagement rather than a shift between locomotion and immobility.

This is consistent with observations of prefrontal activity tracking willingness to work in primates and changes in task utility in rats^45,46^. Here, we cannot distinguish disengagement due to fatigue, satiation, frustration, or a combination of factors, and we therefore operationally describe the neural state preceding disengagement operationally as a loss of “motivation”. For subsequent analyses, we summarized the inferred state by the angular position of the context within the PC1-PC3 plane, providing a scalar neural index of “motivation” (Methods, Fig. 5G).

#### Putative interneurons show biased firing preference for low-motivation states

We used C3PO’s mapping from state to neurons to test if different neuronal subtypes preferentially altered their firing with motivational state. We assigned putative excitatory and inhibitory labels using short-latency peaks and troughs in spike cross-correlograms^47^ (Fig. 5I).

We projected classified neurons’ mark embeddings onto the motivation PCs (Fig. 5J). Putative excitatory neurons spanned embedding angles, whereas putative inhibitory neurons concentrated within a sector (Fig. 5J). This predicts diverse and consistent motivation state preferences among excitatory and inhibitory neurons, respectively. We correlated each neuron’s trial-average firing rate with the inferred motivation state. Excitatory neuron correlations spanned positive and negative values. Putative inhibitory neurons were biased toward negative correlations, consistent with greater activity at lower values of the motivation index (Fig. 5K).

Finally, we ordered neurons with strong projections onto the motivation plane by embedding angle for the two putative classes. Plotting their relative firing rates across run epochs revealed a slow progression of neuronal activation corresponding to the motivation state (Fig. 5L). These observations corroborate the correspondence between the learned embeddings and observed firing, linking the slow population state identified by C3PO to individual neurons and their physiological annotations.

#### Interactions between motivational state and reward responses

We next asked whether motivation state interacts with representations expressed within individual trials. Alongside task-progression signals^40,42^ (Fig. 5D, E), prefrontal populations represent recent reward histories and decision variables relevant to foraging^13,43,44^ (Fig. 6A). These can be represented through nonlinear mixed selectivity^2^, corresponding to neuronal tuning over a joint space of task variables and internal states (Fig. 1A, 6A). We reasoned the slow motivation state could modulate expression of faster task representations, allowing the same event to evoke different population responses based on engagement. To examine this, we focused on the interval between reward outcome and subsequent path choice (Fig. 6B). In the depletion– repletion task, the preceding outcome can inform a subsequent decision to stay in the current patch or switch to a different patch^13,48^, but whether or how motivational state might impact activity was unclear.

**Figure 6.**
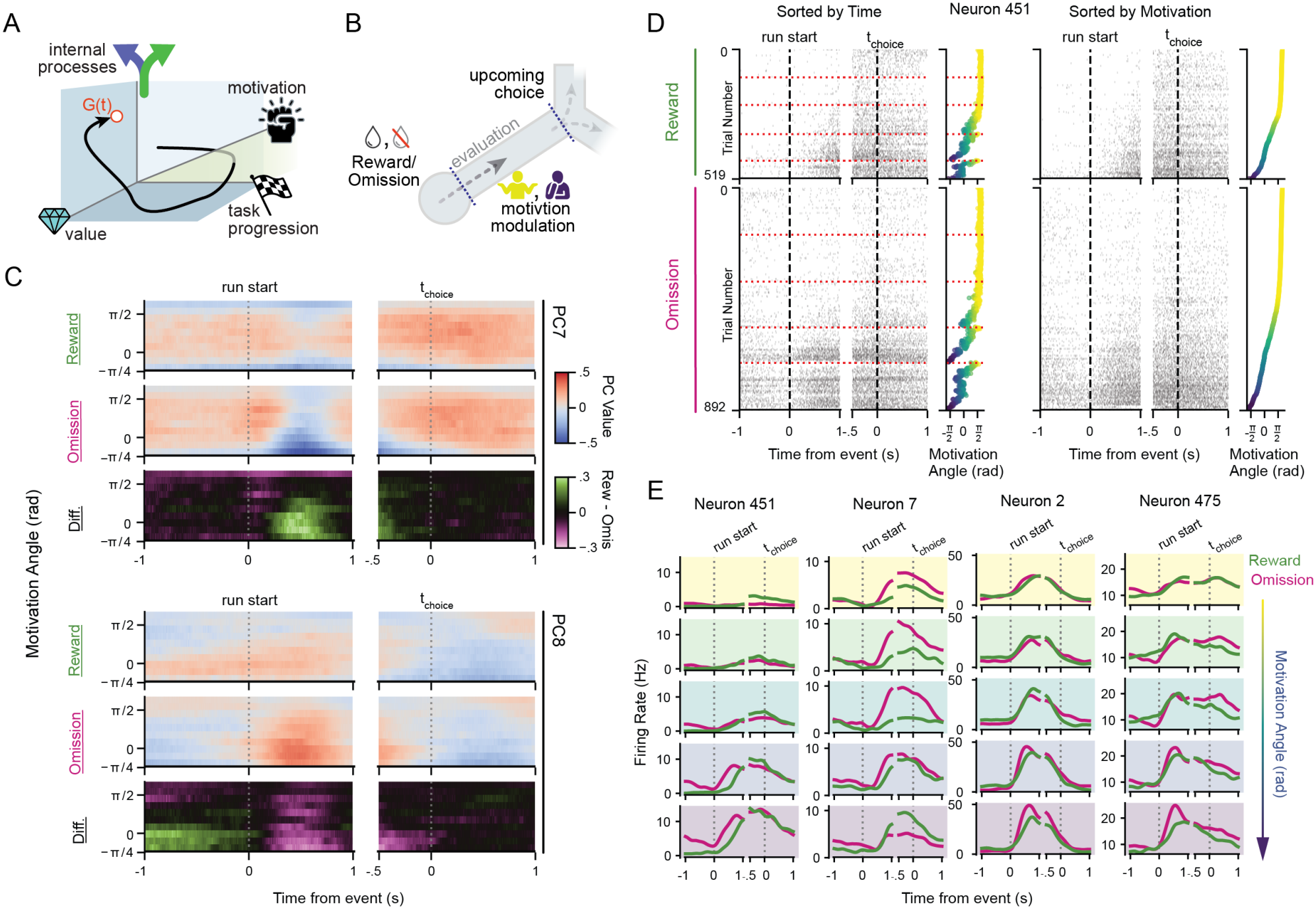
Prefrontal representations of task progression and reward history vary with inferred motivational state. Data are from the rat medial prefrontal cortex recording analyzed in Fig. 5. Trial motivation is indexed by the angular position of trial-averaged C3PO context in the PC1–PC3 plane. **A**, Schematic of candidate generative features underlying prefrontal population activity, including task progression, motivation, value, and other internal processes. **B**, Cartoon illustrating potential interaction between motivational state and the processing of preceding reward delivery or omission during progression toward the next choice. **C**, Mean context values for PC7 (upper block) and PC8 (lower block), grouped by trial motivation and preceding outcome. Within each block, the upper two rows show trials following reward and omission; the bottom row shows their difference (reward minus omission; green, positive; magenta, negative). Separate time windows are aligned to run initiation and choice. Outcome-dependent differences vary with both task phase and motivation, with pronounced differences around run initiation at lower motivation angles. **D**, Spike rasters for an example neuron (neuron 451), separated by preceding reward (top; 519 trials) or omission (bottom; 892 trials). Each row represents one trial and each mark one spike. Trials are ordered chronologically on the left and by decreasing motivation angle on the right. Adjacent scatter plots show the motivation angle of each trial, colored from high (yellow) to low (blue). Red dotted lines mark boundaries between task epochs; vertical dashed lines indicate run initiation and choice. Ordering trials by motivational state reveals changes in firing near run initiation and their dependence on preceding outcome. **E**, Mean firing rates of four example neurons, grouped into five equal-width motivation-angle bins, decreasing from top to bottom, and separated by preceding reward (green) or omission (magenta). Background colors indicate motivation group. Rates were calculated in 50-ms bins and smoothed with a Gaussian kernel (s.d., 100 ms). Response magnitude, timing, and reward–omission preference vary across motivational states. Panels C and E pool non-switch trials across task epochs within the displayed motivation range. In **C–E**, the run-initiation and choice windows are aligned separately; horizontal gaps separate these windows.

We found clear evidence for this sort of interaction. We examined C3PO context over the joint space of task progression, motivation, and recent reward outcome (Fig. 6C). Two components, PCs 7 and 8, showed outcome-dependent temporal profiles that varied across motivation states. Subtracting the omission- and reward-conditioned responses revealed pronounced, temporally structured differences around port departure and run initiation at lower motivation values. Because C3PO’s learned mark embeddings link each context direction to a pattern of neuronal co-modulation (Fig. 1E), the increased separation of reward and omission responses suggests state-dependent modulation of an outcome-sensitive neuronal ensemble. This organization is consistent with a neuronal assembly whose response to reward omission becomes more strongly differentiated around port departure at lower motivation.

We confirmed this modulation is expressed in individual neurons by identifying units with strong embeddings on PCs 7 and 8. For each example neuron, we pooled trials across all run epochs and plotted event-aligned spike rasters separately for trials following reward and omission. Ordering trials by inferred motivation revealed changes in response timing and outcome selectivity more clearly than chronological order (Fig. 6D). Activity emerged around port departure at low motivation, with an earlier rise following omission than reward. We then averaged event-aligned firing rates for additional neurons within motivation bins, separated by preceding outcome (Fig. 6E). These curves revealed state-dependent changes in response magnitude and timing, including stronger omission-associated responses in some neurons and reversals of reward–omission preference in others. These observations corroborate the context-level organization in individual-neuron firing, linking the inferred motivation state to changes in how recent outcomes are represented across trial phases.

### Automated latent variable discovery from multiple datasets

A computational method’s utility depends on ease of use and robustness to free parameter choices. In addition to an extension that makes C3PO easy to use within Spyglass, our reproducible data analysis system^49^, we also developed a command-line interface for applying C3PO directly to recordings in the DANDI archive, a NIH-funded repository for sharing electrophysiological data (Fig. 7A). Users specify the dandiset identifier and path of an NWB asset. The workflow then uses the DANDI API and pyNWB^50^ to locate the file and stream its contents from cloud storage. A C3PO model is then constructed and trained using a predefined architecture and training parameters. The context is then inferred and saved locally with the trained model. We also provide a google-colab compatible notebook for this pipeline to allow users to leverage cloud-compute resources to train models and download results locally for further analysis.

**Figure 7.**
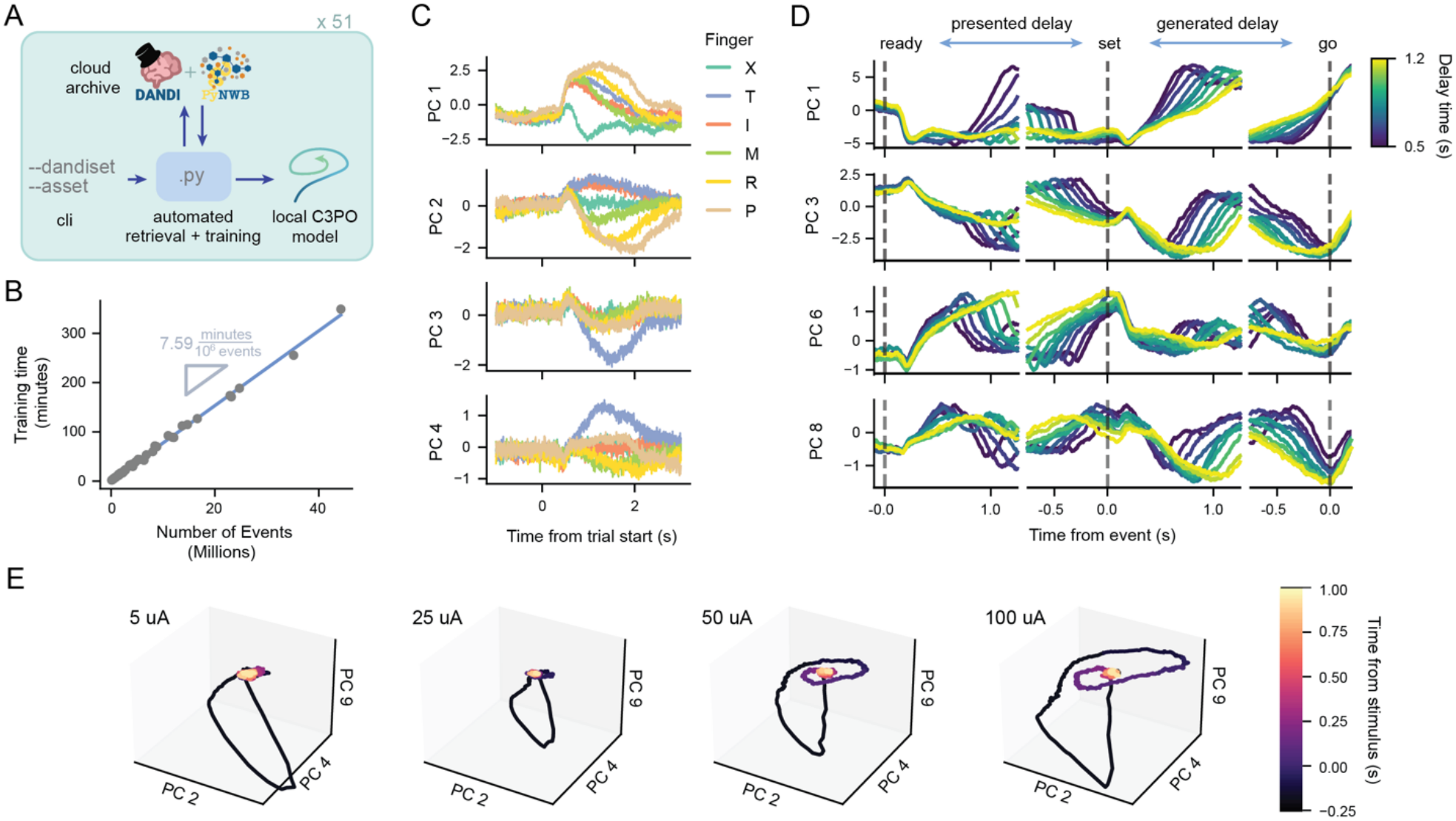
Automated C3PO analysis identifies structured population dynamics across public datasets. **A**, Automated workflow for retrieving recordings from the DANDI archive and training C3PO models. Given a dandiset identifier and Neurodata Without Borders (NWB) asset path, the command-line interface retrieves spike data, trains a model using default parameters and saves the model and inferred representations locally. Archive screening identified 51 candidate recordings containing at least 50 sorted units and 15 min of recording. **B**, Training time versus number of spike events for the 51 recordings, each trained independently on one NVIDIA A100 GPU. Each point represents one dataset; the blue line shows a linear regression, with a slope of 7.59 min per million events. **C**, Human posterior parietal cortex activity during attempted individual finger movements for brain–computer interface control by a participant with tetraplegia^51^. Traces show mean C3PO context PCs 1–4 aligned to trial onset and grouped by instructed movement: thumb (T), index (I), middle (M), ring (R), little finger (P) or no movement (X). Context responses distinguish finger conditions across the trial. **D**, Macaque dorsomedial frontal cortex activity during the Ready–Set–Go interval-reproduction task^52^. The animal estimates the interval between Ready and Set and reproduces it by timing its subsequent movement (Go). Mean context PCs 1, 3, 6 and 8 are shown in separate windows aligned to these three events, indicated by dashed lines. Colors denote the presented interval duration throughout all windows. Context components exhibit interval-dependent ramps and plateaus during both interval estimation and production. **E**, Mouse visual cortex responses to electrical microstimulation^53^. Mean stimulation-aligned trajectories are projected onto context PCs 2, 4, and 9 for the indicated current amplitudes. Each trial’s mean pre-stimulus context, calculated from −250 to 0 ms, was subtracted before averaging. Colors indicate progression through the stimulation-aligned response. Rotational excursions followed by relaxation, particularly at higher amplitudes, are consistent with underdamped population dynamics. All models were trained independently with only spiking activity; task and stimulation annotations were used subsequently to align and group context responses.

To evaluate the utility of this workflow, we undertook a large-scale screen of publicly available neural recordings in the DANDI archive (Fig. 7A). An archive search using GPT-Sol 6.1 identified 51 eligible NWB files from distinct dandisets, each with at least 50 sorted units and 15 minutes of recording data (Table S2). Selected datasets included human, nonhuman primate, and rodent recordings across diverse brain regions, behavioral tasks, sleep states, and experimental perturbations—providing a broad test of C3PO’s applicability across recording scales, experimental conditions, and neural systems.

Each dataset was trained independently using the default protocol on a single NVIDIA A100 GPU. Training time increased linearly with the number of spike events, averaging approximately 7.5 minutes per million events (Fig. 7B). For comparison, acquiring one million events at a population event rate of 2,000 spikes per second takes approximately 8.3 minutes, indicating potential of C3PO as a tool for real-time feedback experiments. Visual inspection of the inferred context representations confirmed non-trivial temporal dynamics in every completed training run, supporting the practical robustness of the default protocol across heterogeneous recordings.

To illustrate the inferred dynamics, we selected three datasets. In human posterior parietal cortex, Guan et al. recorded activity while a tetraplegic participant attempted individual finger movements for brain–computer interface control^51^. C3PO context components showed distinct trial-aligned responses across finger and no-movement conditions (Fig. 7C), consistent with the finger-selective representations reported in the original study.

In macaque dorsomedial frontal cortex recordings during the Ready–Set–Go task, animals estimated and reproduced a visually presented interval^52^. Context components exhibited interval-dependent ramps and plateaus during both estimation and production (Fig. 7D), paralleling the original finding that cortical trajectories evolved toward movement-related states more slowly for longer sample intervals.

Finally, we examined mouse recordings during single-pulse electrical stimulation of visual cortex^53^. The original study identified widespread, amplitude-dependent circuit responses comprising excitation, suppression, and rebound excitation. Stimulation-aligned C3PO context trajectories exhibited rotational excursions followed by relaxation, with post-stimulation loops particularly apparent at higher amplitudes (Fig. 7E). This geometry is consistent with underdamped population dynamics and provides a compact representation of the network response described in the source study.

## DISCUSSION

Our results demonstrate that C3PO recovers interpretable structure associated with the generative features of neural activity across species, brain regions, and recording conditions. Despite the absence of behavioral or physiological annotations during training, the model identified representations related to proprioception, spatial position, head direction, and oscillatory phase, and further linked these population features to single neuron tuning. The learned context preserved dynamics across timescales, from rapidly compressed hippocampal sequences to slow prefrontal states associated with task engagement. It also revealed interactions between these states and trial-specific outcome representations.

C3PO’s performance on sorted spiking datasets was comparable to or better than existing methods^5,8^, generating inferred latents that separated generative features more orthogonally than alternative approaches. In turn, these separated latents enabled simple downstream decoding of the features themselves, with average errors at or below those achieved with other models.

Where C3PO’s utility comes to the forefront is in the architectural features that differentiate it from these models. First, the point-process framework and focus on spike events negates the need for binning or trial segmentation^11^. This avoids a core tradeoff other models^3,7,8,8,54^ face in determining spike bin durations: short millisecond-scale time bins are necessary to track rapid neural dynamics, but become sensitive to spiking stochasticity, resulting in noisier embedding estimates. Instead, C3PO models this stochasticity directly.

Context updates with each detected spike, meaning the temporal resolution depends only on (and dynamically scales with) the event density. In practice this provides access to relevant features spanning many orders of magnitude, from the tens of milliseconds of CA1 replay events to minutes and hours underlying changes in motivation.

Second, C3PO achieves its predictive performance while maintaining a fully interpretable latent geometry that is amenable to numerous downstream analyses. In many methods, the non-linearity of embedding means that distances and linear transformations of the embedded results cannot be rigorously studied for dynamics or their relationship to single neuron activity^7,8,54^. In contrast, C3PO is selective in applying non-linearity: flexible waveform and history encoders capture complex interacting features and temporal dependencies, while the final readout remains linear in the log intensity of spike events. This is conceptually similar to the approach used in latent-dynamical models such as LFADS^5^ and in recent work that attributes inferred latent variables back to neuron rates in other encoding models^16^; however, C3PO’s architecture embeds this interpretability into the geometry of the latent space itself.

Finally, by operating directly on detected events and/or their waveform features, C3PO removes the time and curation required for a separate spike-sorting stage necessary for almost every other unsupervised feature extraction. Paired with the high computational efficiency and hyperparameter robustness demonstrated by our automated pipeline application, this extends the scope of potential C3PO applications to high-throughput experimental designs, real time behavioral feedback, and brain-computer interfaces.

Moreover, both our results and previous works^10,11,37,55^ suggest that rather than obscuring latent features with unfiltered noise, inclusion of un-sortable spike events can enhance feature extraction and decodability. By jointly encoding these unsorted waveforms and sorted spikes in the hybrid architecture, C3PO enables us to use these often-discarded events to improve our understanding of both population and individual neuron activity. Together, these findings show that contrastive prediction of individual spike events provides a common basis for reconstructing established neural representations and identifying hidden population structure, using sorted, clusterless, or hybrid recordings.

C3PO’s point-process modeling also makes it well suited to study how neural representations scale and are modified across different timescales. In CA1, C3PO identifies the common representation of spatial sequences expressed during behavior, theta cycles, and SWR replay despite their compression across timescales. In mPFC, the same framework reveals how outcome representations, which are expressed in ∼500 ms within a relatively small subset of trials, evolve over hours of recording with internal motivation states. This combination of sensitivity, scale, and unsupervised learning uniquely positions C3PO for studying other internally structured neural processes such as sleep and learning.

This work demonstrates the effectiveness of contrastive learning in extracting linearized representations of real-world features from neural spiking activity. This raises the question of whether higher (e.g., cortical) brain regions could plausibly implement similar algorithms to learn representations of external features from input sensory spiking. Interestingly, biologically plausible learning rules consistent with contrastive learning have been developed and shown to be useful^56,57^. Adapting C3PO’s event-based prediction objective to these constraints would allow tests of whether local plasticity can recover similar latent structure, and how neural circuits might generate the comparison signals required for contrastive learning. This would connect predictive representation learning as an analysis method to hypotheses about how neural populations acquire representations of the states underlying their input activity.

## MATERIALS AND METHODS

### C3PO Loss Derivation

#### Conceptual model of neural signals

At each timepoint, the activity of a neural population reflects the accumulated history of internal dynamics, external inputs, and ongoing computation. We define a summary of this high-dimensional sequence of past events as the generative latent variable *G*, which acts as a sufficient statistic of the system history for predicting future activity. Formally, *G* captures all information from the past that is relevant for determining the distribution of future spiking events such that:

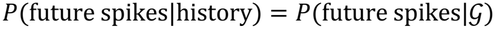

This concept is closely related to the notion of causal states in computational mechanics, where histories are grouped according to their predictive equivalence^58^.

Intuitively, *G* is the (relatively) low dimensional generative feature driving neural activity^1^. For example, while a rat exploring its environment experiences complex sensory inputs, the simple x/y-coordinate position is highly predictive of the spiking probabilities of ‘place cells’ in the hippocampus. More generally, in different brain regions, latent variables may reflect mixtures of environmental, task, or internal states, giving rise to dynamic context-dependent firing patterns.

Within this framework, the probability that a given neuron *i* fires a spike at time *t* is governed by a conditional intensity function: *f*_*i*_(*t*) = *f*_*i*_(*G*_*t*_). In analogy to classical tuning curves, which describe firing rates as functions of experimentally controlled variables, here each neuron exhibits a (generally nonlinear) response function over the latent generative state, reflecting selectivity to one or more underlying variables. Spikes are then generated as stochastic events where a neuron fires spikes probabilistically according to its conditionally independent instantaneous firing rate *f*_*i*_(*G*_*t*_), consistent with standard point-process formulations of neural activity. In this idealized, fully-observed model description, this stochastic sampling reflects the intrinsic entropy of the neural dynamics^58^.

These spike trains and corresponding neuron identities are not observed directly. Rather, implanted extracellular electrodes measure voltage deflections produced by nearby neurons, resulting in observed waveform snippets whose shapes depend on the spatial and biophysical properties of the underlying sources. Following approaches used in spike sorting^9^ and other clusterless-decoding methods^37,59^, each waveform snippet can be reduced in dimensionality and featurized by the peak voltage across each channel, width of the waveform, or other characteristics that define the waveform feature vector *W*_*i*_. Together with the delays between observed spike times, we can represent the resulting data as a sequence of marked point-process observations:

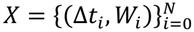

This generative view highlights the central challenge: while we are interested in studying the latent variable *G* that governs neural activity, our observations are a noisy, high-dimensional signal separated from this variable by multiple stochastic processes.

#### Mark Embedding

The first stage of the model constructs a low-dimensional embedding of each observed high-dimensional event waveform 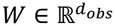. For dense recording devices with hundreds of electrodes, *d*_KMN_∼10^2^. Each waveform is passed through the embedding layer to generate a low-dimensional embedded mark:

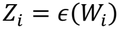

where 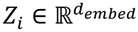 and *d*_*embed*_ ≪ *d*_*obs*_. The embedding function *ϵ* may be implemented as any neural network compatible with backpropagation (e.g., multilayer perceptron or convolutional encoder). This flexibility allows the model to accommodate diverse electrode geometries and recording configurations. Through this work, we use dense three-layer networks for this embedding (Table S1).

#### Context Inference

Waveform embedding transforms our sequence of observations

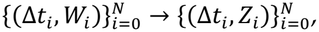

where *i* indexes over spike count starting with the first spike. We next infer a latent context variable *C*_*i*_ that integrates information across past events:

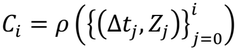

This extracts a learned representation of population firing history that evolves over time (Fig. 1B). Note that this function generalizes the recursive update *C*_*i*_ = *ρ*((Δ*t*_*i*_, *Z*_*i*_), *C*_*i*−1_) and allows the context to be implemented using a wide class of architectures. In general, *ρ* may be any causal sequence model, including recurrent neural networks, temporal convolutional networks, or transformer-based models. Through this work, we use a WaveNet architecture, consisting of stacks of dilated causal convolutions^20^ (Table S1). This architecture is well suited for neural spiking data, as it efficiently captures long-range temporal dependencies while maintaining causal structure. By exponentially expanding the receptive field with depth, dilated convolutions enable the model to integrate information over extended time horizons without the computational cost associated with recurrent or attention-based models^20^.

#### Rate Prediction

We next model the likelihood of future spiking events by specifying the conditional intensity function over the interval *t* ∈ (*t*_*i*−1_, *t*_*i*_]:

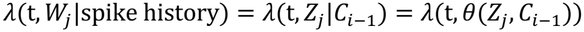

where *Z* and *C* serve as summary statistics for the predictive information contained in a waveform feature and the firing history, respectively. C3PO architecture supports any closed-form hazard function with parameters *θ*. In turn, *θ* is a learned function of the context and embedded mark.

#### Intractability of maximum likelihood

Training a model with continuous waveform features presents a distinct challenge. Namely, the likelihood of any observation (Δ*t*_*i*_, *W*_*i*_) is given by:

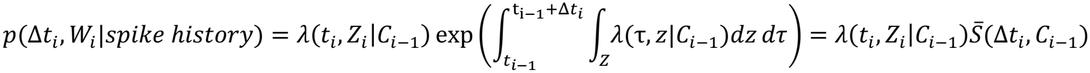

where 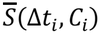 is the probability of no spikes of any waveform type occurring in the interval Δ*t*_*i*_. This can be viewed as analogous to a partition function for these point process likelihoods. Because the embedded marks *Z* exist in a continuous, non-compact space, a closed form solution for this integral cannot be evaluated during training.

#### Contrastive Loss

We therefore introduce an alternative approach derived from Noise Contrastive Estimation (NCE)^22^. Contrastive objectives such as InfoNCE avoid explicitly evaluating a partition function by recasting density estimation as a classification task: identifying an observed sample among a finite set of alternatives drawn from a noise distribution. Classification uses scores proportional to the ratio of the true conditional probability to the noise density. Any normalization factor (e.g., a partition function) shared across candidates cancels when these scores are normalized over the candidate set, eliminating the need to integrate over the full observation space. Here we show a derivation demonstrating this same logic can apply to point process probabilities.

To adapt this to point processes with continuous marks, at each timepoint we sample *n*_*neg*_ events from random alternative locations in the series. We score each mark by the relative probability of it occurring conditioned on the current context:

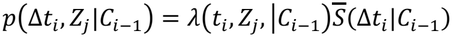

vs. the background, context-independent probability of it occurring (e.g., noisy firing):

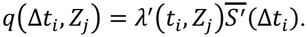

The loss is then a softmax selection of the true next event *Z*_*i*_ from the sampled set based on these scores:

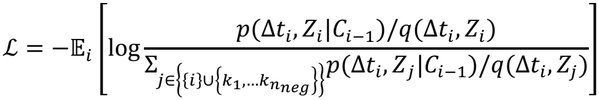

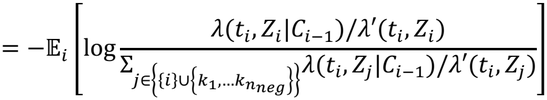

The problematic integrated survival terms cancel in this formulation, as the probability of both positive and negative sampled marks are equally dependent on a lack of other events in the Δ*t*_*i*_ interval. This makes the loss tractable for gradient descent algorithms.

Also, note that optimization maximizes the ratio *λ*(*t, Z*|*C*)/*λ*′(*Z*), which is a description of how much more likely a waveform event is given the context relative to its baseline rate. This in turn maximizes a lower bound for the mutual information between the inferred context and waveform observation^21^. This is a fundamental difference from generative methods that optimize for variance explained in a time series.

#### Choice of conditional process

In high density recordings, total spike event rates are often >1kHz. While more descriptive hazards are available and implemented in the C3PO package, we found that using a Poisson emissions model with constant rate intensities during the <1ms between spike updates was sufficient to capture all features of interest. To further aid geometric interpretation, we implement the intensity as log-linear, giving a hazard function:

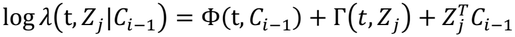

Here, Φ(*t, C*_*i*−1_) is a dynamic mark-independent modulator of firing rate. This term cancels during comparison to negative samples within the contrastive loss function and is not learned by the C3PO model. Similarly, Γ(*t, Z*_*j*_) describes a mark-dependent firing rate coefficient shared by the conditional and background noise models. As this cancels in the ratio score used during training, this is also not explicitly learned within C3PO. We include these terms here both for completeness and to emphasize that C3PO optimizes for information about relative rate changes rather than absolute firing rates.

This formulation places both embedded marks and contexts in a common latent space, where their inner product directly determines the instantaneous event rate for those marks. Intuitively this means that the context is a time-varying vector that sweeps around to point towards the set of embedded waveforms with high conditional firing probability.

#### Choice of noise process

In the simplest case, the noise firing rate is nominally uniform across mark space (*λ*′.(*t, Z*) = Γ(*t, Z*) + *α*). With the choice of conditional hazard above, this gives the loss:

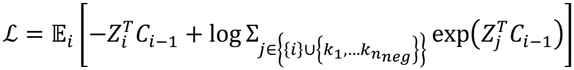

This is exactly the InfoNCE objective used in CPC^21^, establishing a direct correspondence between regularly sampled sequence models and the present continuous-time, point-process setting.

More generally, this reference model can be used to impose structure on the embedding space. We specifically choose a Poisson process that is independent of context, with log intensity decaying with the squared 2-norm of the embedding:

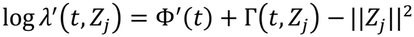

This choice regularizes the mark space by encouraging waveform events that are not predictable from spike history (e.g., noise) to embed near the origin. Empirically, we found using the Euclidean norm stabilizes training and improves convergence. This results in a final loss function:

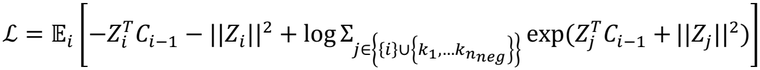

### Model Architecture

#### Waveform Embedding

Spike event data was embedded using either a clusterless, sorted, or hybrid architecture depending on data type (Fig. S1). For clusterless events, the vector of peak amplitudes across recording channels was mapped to the Z embedding space using a fully connected multilayer perceptron with leaky-ReLU activations in the hidden layers and a linear output layer. For multi-shank recording probes, each shank had a separate encoder that processed events originating from its channels, with all encoders producing embeddings in the shared Z space. Network weights were initialized using He-uniform initialization; layer widths and embedding dimensionalities are provided in Supplementary Table 1. For sorted spikes, each event’s categorical neuron label was mapped to a learned, neuron-specific mark embedding through a lookup table, equivalent to a linear projection of a one-hot neuron-identity vector. Embedding weights were initialized from a zero-mean Gaussian distribution with a standard deviation of 0.1; embedding dimensionalities are provided in Supplementary Table 1. The hybrid encoder routed each spike event to one of two branches according to its sorting assignment: spikes that were assigned to a single unit during sorting were represented by learned neuron-specific embeddings, whereas unassigned events were embedded from their waveform features using a multilayer perceptron. Both branches produced embeddings in the same latent mark space, allowing a shared context network to integrate the combined event history, with both encoders and the context network optimized jointly under the contrastive objective.

#### Context Embedding

Context embeddings were computed using a WaveNet-based network of dilated causal convolutions with gated activations and residual and skip connections^20^. Each input comprised the event’s mark embedding concatenated with an additional variable corresponding to the logarithm of its preceding inter-event interval, with convolutions operating over the ordered event sequence. Layer configurations and context dimensionalities are provided in Supplementary Table 1.

### Spike Sorting

Sorted spike ids and event times were taken from published data in the original publications^13,23,24,28^.

### Waveform extraction

Clusterless waveform features were extracted from the same filtered recording used for spike sorting. Event times were identified using DetectPeakByChannel in the spikeinterface package, with an amplitude threshold of 100 µV and a local exclusion radius of 100 µm. Waveforms of each event were featurized by the peak amplitude within ±500 µs of the event on each channel of the tetrode on which the event was detected. To improve numerical conditioning before training, hippocampal tetrode waveform-amplitude vectors were centered by subtracting each event’s mean amplitude across channels and then divided by a fixed factor of 1,000. For PFC probe waveform data, each event’s amplitude vector was divided by its maximum absolute amplitude across channels and then centered by subtracting the mean of the rescaled vector. After normalization, the magnitude of noise in channels far from the event preserves information about the event amplitude for the encoder neural network.

### Data preparation

Events from all included neurons or recording shanks were merged and sorted chronologically, and inter-event intervals were calculated from successive timestamps, expressed in milliseconds. Coincident timestamps were handled by removing zero-interval events. Alternatively, the timing of coincident events can be numerically jittered by a small *ϵ*∼*U*(−.1 *ms*, .1 *ms*) to desynchronize them for training, though this approach is not used in any of the models reported here. Event–interval pairs were segmented into overlapping sequences containing a fixed number of events.

### Model Training

The waveform encoder, context network, and hazard-prediction model were trained jointly by minimizing the contrastive loss using the Adam optimizer. Gradients were computed by automatic differentiation in JAX, with just-in-time compilation used to accelerate training. Training sequences were randomly shuffled at the beginning of each epoch and processed in minibatches; incomplete final minibatches were excluded. For each prediction, negative examples were sampled randomly with replacement from event embeddings within the current minibatch. The mean loss across minibatches was recorded after each epoch to monitor convergence and guide the adaptive hyperparameter schedule described below. All models were trained on a single A100 GPU, except for the prefrontal data, which was trained using 4 A100 devices using multi-device optimizations implemented in the c3po package.

Training used an adaptive schedule for negative sampling and minibatch size while maintaining a fixed learning rate. After a minimum waiting period following initialization for each adjustment (five epochs by default), training was considered stalled when the mean loss over the preceding four epochs was less than 1.01 times the current epoch’s loss. At each stall, the number of negative samples was doubled up to a specified maximum; once this maximum was reached, subsequent stalls instead halved the minibatch size, subject to a specified minimum, and reset the negative-sample count to the larger of its initial value and one-eighth of its maximum. This schedule continued until training stalled at both the maximum negative-sample count and minimum minibatch size, or the maximum number of epochs was reached.

### Synthetic data generation

Simulated marked spike trains were generated for 64 neurons over 5,000 s, driven by a latent phase that advanced uniformly with a period of 3 s. Each neuron initially had a Gaussian phase-tuning curve defined using the shortest circular distance from its preferred phase. Preferred phases were sampled uniformly across the full cycle, tuning widths uniformly from 0.1–0.5 radians, and peak firing rates uniformly from 3– 10 Hz. Spike events were generated by drawing exponential waiting times using the summed population firing rate and selecting the emitting neuron with probability proportional to its firing rate. Rates were recalculated after each event or after 0.3 s without an event. Each neuron was assigned a fixed, 128-channel waveform-feature template consisting of a Gaussian amplitude profile across channels, with a uniformly sampled center channel, peak amplitude of 0.5–1.5 arbitrary units, and standard deviation of 0.5–3 channels. Template parameters were sampled independently of phase-tuning parameters. Individual spike marks were generated by adding independent, zero-mean Gaussian noise with a standard deviation of 0.1 to each channel of the corresponding template.

To introduce “noise” neurons without latent-phase tuning, the spike times of 16 neurons were independently reassigned to uniformly sampled times throughout the recording, preserving their spike counts and associated waveform features and leaving 48 phase-tuned neurons. All events were sorted chronologically, and their waveform-feature vectors and inter-spike intervals used as inputs to C3PO; neuron identities and latent phases were retained for evaluation.

### Analysis Tools

Post-training analyses were performed using the ‘C3poAnalysis’ class in the c3po package, which provides a common interface for applying trained models and characterizing their learned representations. Each analysis object stores the model configuration and parameters alongside mark embeddings, context trajectories, and their timestamps, and maintains derived representations such as interpolated trajectories and principal-component scores. The class provides methods for feature-conditioned summaries, event alignment, spectral and correlation analyses, and supervised decoding of behavioral or physiological variables from context. Model configurations, trained parameters, and event-time embeddings can be saved and reloaded for reuse across analyses. Details of methods used in this paper are given below.

#### Post Training Feature Inference

Trained models were applied to spiking timeseries using the ‘embed_data’ method. Long recordings were embedded in successive chunks (50,000 events by default), each preceded by 2,000 events of overlapping history; outputs corresponding to this overlap were discarded before concatenation, and the final incomplete chunk was retained. The first 2,000 events of each input series were marked as missing to exclude contexts lacking the prescribed history. For analyses requiring evenly spaced timestamps, context trajectories were linearly interpolated onto regular time steps. PCA was fitted to non-missing event-time contexts within the selected analysis intervals, with the resulting transformation applied to both event-time and interpolated contexts. Mark embeddings (Z) were projected onto the principal axes fitted to the context embeddings, without mean subtraction, yielding event- or neuron-specific coefficients describing their loading on each context component.

#### Feature Modulated activity

For analysis of the relationship between a predicted feature (e.g., theta phase, position, orientation) and the inferred context we used the ‘bin_context_by_feature’ and ‘bin_context_by_feature_2d’ methods. Context vectors or their principal-component scores were restricted to valid analysis intervals and grouped into bins of a single behavioral or physiological feature, or joint bins of two features, using timestamp-based alignment. Bins were defined by specified edges or uniformly spaced edges spanning the observed feature ranges and retained the associated context samples for subsequent calculation of feature-conditioned response summaries.

#### Time Aligned Responses

The ‘aligned_response’ method in the c3po package was used to characterize event-related changes in inferred context by extracting windows of interpolated context vectors around specified event times. Each event was aligned to the sample at, or immediately preceding, its timestamp. Windows extending beyond the available data or containing missing context values were excluded, and the remaining trajectories were retained individually for subsequent calculation of event-aligned response summaries.

#### Power spectra

Power spectral densities of interpolated context vectors were estimated using the ‘power_spectrum’ method, which applies Welch’s method with Hamming windows and 50% overlap. Default settings used 1,000-sample windows and a 10,000-point Fourier transform, and intervals shorter than one window were excluded. When multiple intervals were analyzed, spectra were summarized at each frequency by the median and 25th–75th percentiles across intervals, with each interval weighted equally.

### Supervised Feature Decoding

Decoding of behavioral and physiological variables from inferred context was implemented through the ‘initialize_decoder’, ‘fit_decoder’, and ‘predict_decoder’ methods of ‘C3poAnalysis’.

#### Macaque Reach Task

Continuous behavioral and biomechanical variables were decoded using the training and validation trial assignments supplied in the NWB file^60^, restricting analysis to active reaches from 100 ms before to 500 ms after movement onset. Context vectors were linearly interpolated at 1-ms resolution and smoothed using Gaussian kernels with standard deviations of 100 ms for the hand-position examples (Fig. S3D) and 40 ms for the comparison across proprioceptive features (Fig. S3F). A multi-output k-nearest-neighbor regressor with 300 neighbors, Euclidean distance, and inverse-distance weighting was fitted using all context dimensions to predict the recorded hand, joint, and muscle-derived variables. Training trials were randomly subsampled without replacement to equalize the number of trials across the eight reach directions, using the least represented direction to determine the count. Predictions were evaluated by pooling time samples across validation trials and calculating the squared Pearson correlation between predicted and observed values for each feature; the 25 highest-scoring features were displayed in Fig. S3F.

Categorical movement information was decoded from a single representation of each trial, obtained by averaging the context principal-component scores from 100 ms before to 500 ms after movement onset. Multinomial logistic regression was used to classify the eight reach directions on active trials, and binary logistic regression was used to distinguish active reaches from passive perturbations, with a maximum of 200 optimization iterations for each classifier. Classifiers were fitted using dataset-defined training trials and evaluated on retained validation trials. Performance was quantified as the fraction of correctly classified trials, and the reach-direction confusion matrix was normalized within each true direction. Both C3PO representation and PCA transformation were held fixed during supervised decoder fitting and evaluation.

#### Rat MEC Head Direction

Head direction was decoded from all 16 principal components of the C3PO context, linearly interpolated at 1-ms intervals, using multinomial logistic regression and circular k-nearest-neighbor regression. Logistic regression used 100 uniformly spaced angular bins with inverse-frequency class weights, and decoded direction was calculated as the posterior-weighted circular mean of bin centers. The nearest-neighbor decoder predicted the sine and cosine of head direction by averaging across 30 uniformly weighted Euclidean neighbors and converted the resulting vector to an angle. Both decoders used five-fold temporal cross-validation. Signed decoding error was calculated as the shortest angular difference between predicted and measured head direction, and predictions were compared with the internal-direction estimates supplied with the original dataset^24^.

#### Rat Hippocampal Recording

Linearized position was decoded from the 20-dimensional C3PO context using multinomial logistic regression, yielding probabilities over 0.5-cm spatial bins. Training targets were Gaussian-smoothed across position bins with a standard deviation of 4 cm, and observations were weighted inversely by spatial-bin occupancy. Contexts were Gaussian-smoothed with a standard deviation of five events, and each position measurement was paired with the context at or immediately preceding its timestamp. Decoder fitting used 100 Adam updates with a learning rate of 0.01 and an L2 regularization coefficient of 10^−^. Predictions were generated using five-fold temporal cross-validation. Point decoding estimates were calculated as posterior-weighted means of spatial-bin centers.

### Inferred tuning curves from C3PO embedding

To infer tuning curves for simulated neurons (Fig. 1) we used the following approach. For each phase-modulated simulated neuron, waveform embeddings were averaged across its spikes using the known neuron labels, and its inferred tuning curve was calculated by exponentiating the dot product of this mean embedding with the mean context vector in each latent-phase bin. Inferred curves were normalized by their mean across phase bins and displayed alongside the ground-truth tuning curves, with neurons ordered by their ground-truth preferred phase (Fig. 1J).

### Firing rate analysis

#### MEC head direction tuning

Head-direction tuning curves were calculated by assigning each firing-rate sample to the head-direction measurement at or immediately preceding its timestamp and averaging rates within uniformly spaced angular bins spanning the full circle. Each neuron’s tuning curve was normalized by its mean across angular bins.

#### mPFC

For event-aligned responses (Figs. 5E and 6E), peri-event spike histograms with were calculated for each unit using 2 ms bins. Counts were normalized to obtain firing rates in Hz, then Gaussian-smoothed with a 15 ms standard deviation along the time axis. For Fig. 5E, rates were displayed as log_2_ ratios to each neuron’s baseline, calculated as the mean across all windows.

To characterize slower activity changes (Fig. 5L), firing rates were calculated in overlapping 60 s windows across all run epochs. Rates were normalized by each neuron’s mean across all analyzed windows.

### Putative excitatory/inhibitory cell type assignment

Putative excitatory and inhibitory cell types were identified from pairwise spike cross-correlograms using custom Spyglass tables. Cross-correlograms were computed in 0.5 ms bin widths and normalized into firing rates. Surrogate correlograms were generated by independently jittering each spike in both trains using uniformly distributed offsets in a 5ms window and applying the same correlation processing. The minimum and maximum surrogate values at each lag from 1000 such shuffles defined lower and upper envelopes.

Interactions were classified as excitatory or inhibitory when the observed correlogram averaged over 1-4 ms lags exceeded the corresponding envelope value. In our analyses, we exclude units that received both excitatory and inhibitory labels from this procedure.

### Motivation state

In the rat prefrontal data, the motivation state at each timepoint was calculated as *M*_*t*_ = atan2(*C*_*pc1,t*_ –*C*_*pc3,t*_). The ordering and negative term in the PCs is arbitrary and chosen such that the score traversed from high to low values through the day. Per-trial motivation values were determined by averaging this score from 500 ms before and after the run start and port poke, respectively.

### Alternative model training

CEBRA^8^ and LFADS^5^ were applied to the same MEC recording used for C3PO. Spike times from all 460 neurons were converted to population spike-count vectors in 8.33-ms bins without additional smoothing. Both models produced 16-dimensional representations, and neither received behavioral or physiological annotations during training.

CEBRA (version 0.6.1) used a multilayer perceptron encoder (’offset1-model’) that mapped each bin independently to a latent vector. Positive pairs were selected using temporal sample indices with a nominal offset of two bins (16.7 ms). Training used a cosine-similarity contrastive objective with a fixed temperature of 1 and the Adam optimizer, with a learning rate of 0.0003, minibatches of 512 samples, and 10,000 optimization steps.

LFADS used the lfads-torch implementation and its bundled ‘nlb_mc_maze’ configuration, comprising a recurrent generator, inferred inputs, and a Poisson observation model. Counts were divided into nonoverlapping, 1-s sequences within continuous recording intervals; incomplete sequences were excluded. The first 80% of eligible intervals were used for training and the remaining 20% for validation. Training used Adam with an initial learning rate of 0.004 and minibatches of 64 sequences for 300 epochs. Factor trajectories were averaged across 50 posterior samples. Both methods returned one latent vector per retained time bin, and PCA was applied separately to each representation for comparison with the C3PO context components.

## ACKNOWLEDGEMENTS

We thank Vanessa Bender for comments and edits on the manuscript. Claude and chatGPT were used to suggest phrasing and edits to the manuscript.

## AUTHOR CONTRIBUTIONS

S.R.B conceived the study. S.R.B developed the statistical framework, model architecture, and software implementation of C3PO. S.R.B, E.L.D, U.T.E., L.M.F., and A.D.K. contributed to interpretation of C3PO features. A.E.C. and A.J. contributed experimental data, and S.R.B., K.K., A.E.C., A.J., and X.S. curated the datasets. S.R.B and C.J.L. designed and performed the analyses, and S.R.B. prepared the figures. L.M.F., U.T.E., and A.D.K. secured funding. S.R.B and L.M.F wrote the manuscript with input from all other authors.

## DATA AND CODE AVAILABILITY

The C3PO software package developed in this study is openly available on GitHub (https://github.com/samuelbray32/c3po) and distributed through the Python Package Index (PyPI) as c3po-neuro (https://pypi.org/project/c3po-neuro/). A cloud-based Google Colab notebook (https://colab.research.google.com/github/samuelbray32/c3po/blob/main/tutorial/c3po_dandi_colab.ipynb) enables users to run C3PO using cloud computational resources without local installation, with options to save results to Google Drive or download them locally. Additionally, an extension integrating C3PO into the Spyglass neural data analysis framework is available at https://github.com/samuelbray32/spyglass_c3po, with a PyPI release as spyglass_c3po (https://pypi.org/project/spyglass-c3po).

Custom analysis code used to generate the results and figures presented in this study is available at https://github.com/samuelbray32/c3po_publication. This repository also provides instructions for downloading and running Docker containers containing the pre-installed software, data, and SQL databases required for complete analysis reproduction.

Previously published macaque primary somatosensory cortex recordings^23^ are available from the DANDI Archive (Dandiset 000127^61^) through the Neural Latents Benchmark project^60^. Rat medial entorhinal cortex recordings are available through EBRAINS^62^ (recording ‘27765_2.mat’)^24,62^. Rat medial prefrontal cortex recordings during foraging and hippocampal recordings during linear-track behavior are available from the DANDI Archive (002044, doi pending publication). The three example datasets plotted in Fig. 7—human posterior parietal cortex recordings during imagined finger movements^51^, macaque dorsomedial frontal cortex recordings during interval reproduction^52^, and mouse visual cortex stimulations^53^—are available from the DANDI Archive under accession numbers 000147^63^, 000130^64^, and 000776^65^, respectively. All additional datasets analyzed in Fig. 7 are available through DANDI; dataset identifiers, file paths, and original publication references are listed in Supplementary Table S2.

Trained C3PO models are available from the Hugging Face dataset repository amuelbray32/c3po_publication (https://huggingface.co/datasets/samuelbray32/c3po-publication, doi pending publication) and can be accessed using tooling in the c3po python package. Dataset-specific loading code and code for generating the synthetic data are provided in the publication repository.

## REFERENCES

1. Vyas, S., Golub, M. D., Sussillo, D. & Shenoy, K. V. Computation Through Neural Population Dynamics. Annu. Rev. Neurosci. 43, 249–275 (2020).

2. Rigotti, M. et al. The importance of mixed selectivity in complex cognitive tasks. Nature 497, 585–590 (2013).

3. Yu, B. M. et al. Gaussian-Process Factor Analysis for Low-Dimensional Single-Trial Analysis of Neural Population Activity. J. Neurophysiol. 102, 614–635 (2009).

4. Churchland, M. M. et al. Neural population dynamics during reaching. Nature 487, 51–56 (2012).

5. Pandarinath, C. et al. Inferring single-trial neural population dynamics using sequential auto-encoders. Nat. Methods 15, 805–815 (2018).

6. Abbaspourazad, H., Erturk, E., Pesaran, B. & Shanechi, M. M. Dynamical flexible inference of nonlinear latent factors and structures in neural population activity. Nat. Biomed. Eng. 8, 85–108 (2024).

7. Gosztolai, A., Peach, R. L., Arnaudon, A., Barahona, M. & Vandergheynst, P. MARBLE: interpretable representations of neural population dynamics using geometric deep learning. Nat. Methods 22, 612–620 (2025).

8. Schneider, S., Lee, J. H. & Mathis, M. W. Learnable latent embeddings for joint behavioural and neural analysis. Nature 617, 360–368 (2023).

9. Buccino, A. P., Garcia, S. & Yger, P. Spike sorting: new trends and challenges of the era of high-density probes. Prog. Biomed. Eng. 4, 022005 (2022).

10. Kloosterman, F., Layton, S. P., Chen, Z. & Wilson, M. A. Bayesian Decoding using Unsorted Spikes in the Rat Hippocampus. J Neurophysiol https://doi.org/10.1152/jn.01046.2012 (2013) doi:10.1152/jn.01046.2012.

11. Deng, X., Liu, D. F., Kay, K., Frank, L. M. & Eden, U. T. Decoding position from multiunit activity using a marked point process filter. BMC Neurosci. 16, 1–1 (2015).

12. Coulter, M. E. et al. Closed-loop modulation of remote hippocampal representations with neurofeedback. Neuron S0896627324009231 (2025) doi:10.1016/j.neuron.2024.12.023.

13. Sun, X. et al. Meta-learning is expressed through altered prefrontal cortical dynamics. Preprint at 10.64898/2026.01.01.697272 (2026).

14. Williams, A. H. et al. Unsupervised Discovery of Demixed, Low-Dimensional Neural Dynamics across Multiple Timescales through Tensor Component Analysis. Neuron 98, 1099–1115.e8 (2018).

15. Williams, A. H. et al. Discovering Precise Temporal Patterns in Large-Scale Neural Recordings through Robust and Interpretable Time Warping. Neuron 105, 246–259.e8 (2020).

16. Schneider, S., Laiz, R. G., Filippova, A., Frey, M. & Mathis, M. W. Time-series attribution maps with regularized contrastive learning. https://doi.org/10.48550/ARXIV.2502.12977 (2025) doi:10.48550/ARXIV.2502.12977.

17. Shchur, O., Türkmen, A. C., Januschowski, T. & Günnemann, S. Neural Temporal Point Processes: A Review. Preprint at 10.48550/arXiv.2104.03528 (2021).

18. Guo, R., Li, J. & Liu, H. INITIATOR: Noise-contrastive Estimation for Marked Temporal Point Process. in Proceedings of the Twenty-Seventh International Joint Conference on Artificial Intelligence 2191–2197 (International Joint Conferences on Artificial Intelligence Organization, Stockholm, Sweden, 2018). doi:10.24963/ijcai.2018/303.

19. Shibue, R. & Iwata, T. Marked point process variational autoencoder with applications to unsorted spiking activities. PLOS Comput. Biol. 20, e1012620 (2024).

20. Oord, A. van den et al. WaveNet: A Generative Model for Raw Audio. Preprint at 10.48550/ARXIV.1609.03499 (2016).

21. Oord, A. van den, Li, Y. & Vinyals, O. Representation Learning with Contrastive Predictive Coding. Preprint at 10.48550/arXiv.1807.03748 (2019).

22. Gutmann, M. U. & Hyvärinen, A. Noise-Contrastive Estimation of Unnormalized Statistical Models, with Applications to Natural Image Statistics. J. Mach. Learn. Res. 13, 307–361 (2012).

23. Chowdhury, R. H., Glaser, J. I. & Miller, L. E. Area 2 of primary somatosensory cortex encodes kinematics of the whole arm. eLife 9, e48198 (2020).

24. Vollan, A. Z., Gardner, R. J.Moser, M.-B. & Moser, E. I. Left–right-alternating theta sweeps in entorhinal–hippocampal maps of space. Nature 639, 995–1005 (2025).

25. Mitchell, S. J. & Ranck, J. B. Generation of theta rhythm in medial entorhinal cortex of freely moving rats. Brain Res. 189, 49–66 (1980).

26. Moser, E. I. et al. Grid cells and cortical representation. Nat. Rev. Neurosci. 15, 466–481 (2014).

27. Sargolini, F. et al. Conjunctive Representation of Position, Direction, and Velocity in Entorhinal Cortex. Science 312, 758–762 (2006).

28. Disruption of theta-timescale spiking impairs learning but spares hippocampal replay. Preprint at 10.7554/eLife.109375.2 (2026).

29. Mizuseki, K., Sirota, A., Pastalkova, E. & Buzsáki, G. Theta Oscillations Provide Temporal Windows for Local Circuit Computation in the Entorhinal-Hippocampal Loop. Neuron 64, 267–280 (2009).

30. Somogyi, P. & Klausberger, T. Defined types of cortical interneurone structure space and spike timing in the hippocampus. J. Physiol. 562, 9–26 (2005).

31. O’Keefe, J. & Recce, M. L. Phase relationship between hippocampal place units and the EEG theta rhythm. Hippocampus 3, 317–330 (1993).

32. Skaggs, W. E., McNaughton, B. L., Wilson, M. A. & Barnes, C. A. Theta phase precession in hippocampal neuronal populations and the compression of temporal sequences. Hippocampus 6, 149–172 (1996).

33. Foster, D. J. & Wilson, M. A. Hippocampal theta sequences. Hippocampus 17, 1093–1099 (2007).

34. Johnson, A. & Redish, A. D. Neural Ensembles in CA3 Transiently Encode Paths Forward of the Animal at a Decision Point. J. Neurosci. 27, 12176–12189 (2007).

35. Firing rate adaptation affords place cell theta sweeps, phase precession, and procession. eLife 12, RP87055 (2024).

36. Klausberger, T. & Somogyi, P. Neuronal Diversity and Temporal Dynamics: The Unity of Hippocampal Circuit Operations. Science 321, 53–57 (2008).

37. Denovellis, E. L. et al. Hippocampal replay of experience at real-world speeds. eLife 10, e64505 (2021).

38. Wilson, M. A. & McNaughton, B. L. Reactivation of Hippocampal Ensemble Memories During Sleep. Science 265, 676–679 (1994).

39. Buzsáki, G. Hippocampal sharp wave-ripple: A cognitive biomarker for episodic memory and planning. Hippocampus 25, 1073–1188 (2015).

40. Muysers, H. et al. A persistent prefrontal reference frame across time and task rules. Nat. Commun. 15, 2115 (2024).

41. Muysers, H., Bartos, M. & Sauer, J.-F. Conjoint generalized and trajectory-specific coding of task structure by prefrontal neurons. Cell Rep. 44, 115420 (2025).

42. Guidera, J. A. et al. Regional specialization in prefrontal cortex manifests in the reliability of task progression codes. Neuron 114, 1152–1161.e8 (2026).

43. Bukwich, M. et al. Competitive integration of time and reward explains value-sensitive foraging decisions and frontal cortex ramping dynamics. Neuron 113, 3458–3475.e12 (2025).

44. Lopez-Yepez, J. S. et al. Time-dependent deployment of medial prefrontal cortical representations in male mice. Nat. Commun. 17, 1473 (2026).

45. Porter, B. S. & Hillman, K. L. Dorsomedial prefrontal neural ensembles reflect changes in task utility that culminate in task quitting. J. Neurophysiol. 126, 313–329 (2021).

46. San-Galli, A., Varazzani, C., Abitbol, R., Pessiglione, M. & Bouret, S. Primate Ventromedial Prefrontal Cortex Neurons Continuously Encode the Willingness to Engage in Reward-Directed Behavior. Cereb. Cortex 28, 73–89 (2018).

47. Fujisawa, S., Amarasingham, A., Harrison, M. T. & Buzsáki, G. Behavior-dependent short-term assembly dynamics in the medial prefrontal cortex. Nat. Neurosci. 11, 823–833 (2008).

48. Comrie, A. E. et al. Hippocampal representations of alternative possibilities are flexibly generated to meet cognitive demands. Neuron S0896627326006744 (2026) doi:10.1016/j.neuron.2026.08.023.

49. Lee, K. H. et al. Spyglass: a framework for reproducible and shareable neuroscience research. eLife 14, (2026).

50. Rübel, O. et al. The Neurodata Without Borders ecosystem for neurophysiological data science. eLife 11, e78362 (2022).

51. Guan, C. et al. Stability of motor representations after paralysis. eLife 11, e74478 (2022).

52. Sohn, H., Narain, D., Meirhaeghe, N. & Jazayeri, M. Bayesian Computation through Cortical Latent Dynamics. Neuron 103, 934–947.e5 (2019).

53. Hickman, J. L., Hughes, G., Sahai, E., Miles, M. & Denman, D. J. Neural population dynamics of direct electrical stimulation of neocortex. Cell Rep. 45, 117420 (2026).

54. McInnes, L., Healy, J. & Melville, J. UMAP: Uniform Manifold Approximation and Projection for Dimension Reduction. Preprint at 10.48550/arXiv.1802.03426 (2020).

55. Trautmann, E. M. et al. Accurate Estimation of Neural Population Dynamics without Spike Sorting. Neuron 103, 292–308.e4 (2019).

56. Bellec, G. et al. A solution to the learning dilemma for recurrent networks of spiking neurons. Nat. Commun. 11, 3625 (2020).

57. Illing, B., Ventura, J., Bellec, G. & Gerstner, W. Local plasticity rules can learn deep representations using self-supervised contrastive predictions. Preprint at 10.48550/ARXIV.2010.08262 (2020).

58. Crutchfield, J. P., Ellison, C. J. & Mahoney, J. R. Time’s Barbed Arrow: Irreversibility, Crypticity, and Stored Information. Phys. Rev. Lett. 103, 094101 (2009).

59. Zhang, Y. et al. Bypassing spike sorting: Density-based decoding using spike localization from dense multielectrode probes. Preprint at 10.1101/2023.09.21.558869 (2023).

60. Pei, F. et al. Neural Latents Benchmark ‘21: Evaluating latent variable models of neural population activity. Preprint at 10.48550/arXiv.2109.04463 (2022).

61. Miller, L. Area2_Bump: macaque somatosensory area 2 spiking activity during reaching with perturbations. DANDI Archive 10.48324/DANDI.000127/0.220113.0359.

62. Vollan, A. Z., Gardner, R.Moser, M.-B. & Moser, E. Left-right-alternating theta sweeps in the entorhinal-hippocampal spatial map (v1). EBRAINS 10.25493/R5FR-EDG (2024).

63. Guan, C., Aflalo, T., Zhang, C. & Andersen, R. PPC_Finger: human posterior parietal cortex recordings during attempted finger movements. DANDI Archive 10.48324/DANDI.000147/0.221122.2256.

64. Sohn, H. & Jazayeri, M. DMFC_RSG: macaque dorsomedial frontal cortex spiking activity during time interval reproduction task. DANDI Archive 10.48324/DANDI.000130/0.241017.1448.

65. Denman, D. & Hickman, J. Spatial and temporal propagation of spiking activity in response to electrical stimulation in visual cortex. DANDI Archive 10.48324/DANDI.000774/0.260520.1753.

