## Supplemental Figures for "Learning predictive latent structure from unclustered neural spike trains"

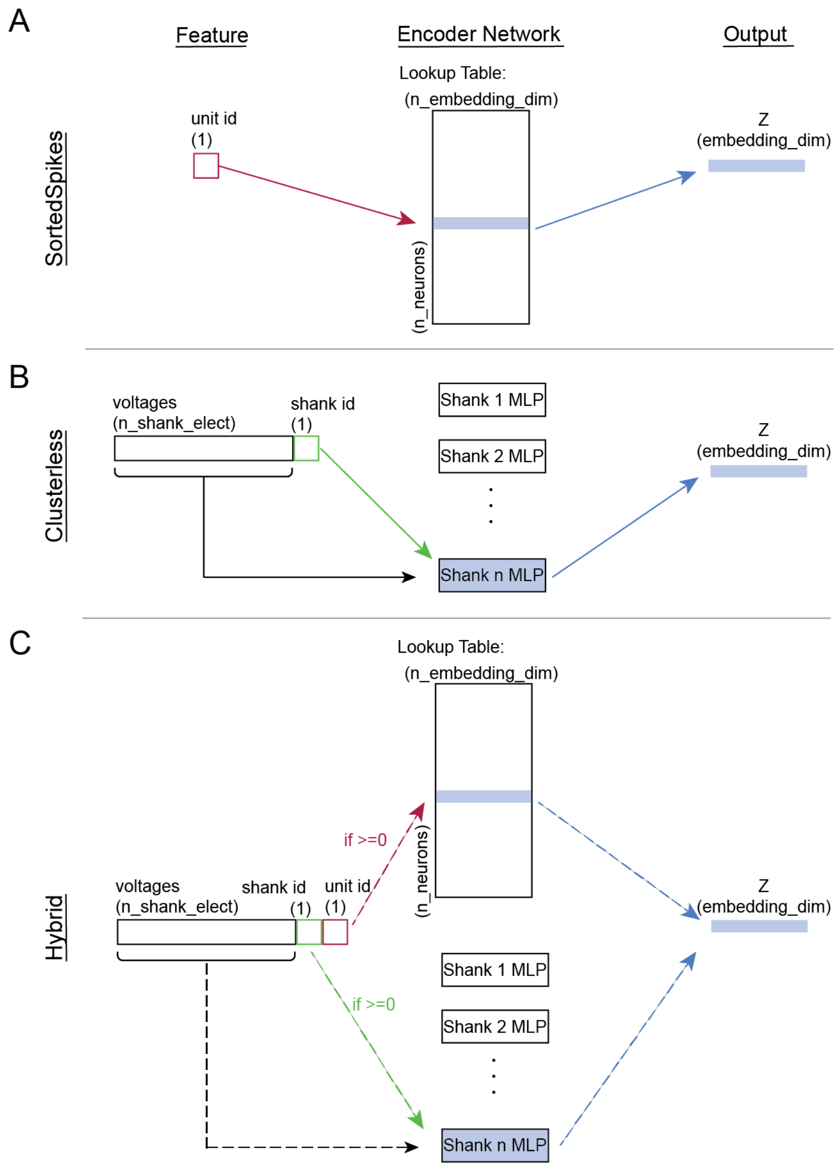


**Supplementary Figure 1.** | **Mark encoder architectures for sorted, clusterless, and hybrid spike inputs.** Each encoder maps individual observed marks (W) into a learned embedding space (Z).

**A,** Sorted-spike encoder. Unit identity, represented as a one-hot vector or integer index, selects a unit-specific embedding from a trainable matrix.

**B,** Clusterless encoder. Continuous waveform features are transformed into embeddings by a multilayer perceptron with leaky rectified linear activations in the hidden layers and a linear output layer. For recordings containing multiple electrode groups, such as probe shanks or tetrodes, group labels route events to separate encoders whose outputs occupy a common embedding space.

**C,** Hybrid encoder. Events assigned to sorted units are embedded using unit identity, whereas unassigned events are embedded using their waveform features. Both branches produce embeddings of the same dimensionality, which are assembled in their original temporal order into a single event sequence for the context network. This architecture incorporates identified units and remaining clusterless events within one C3PO model.

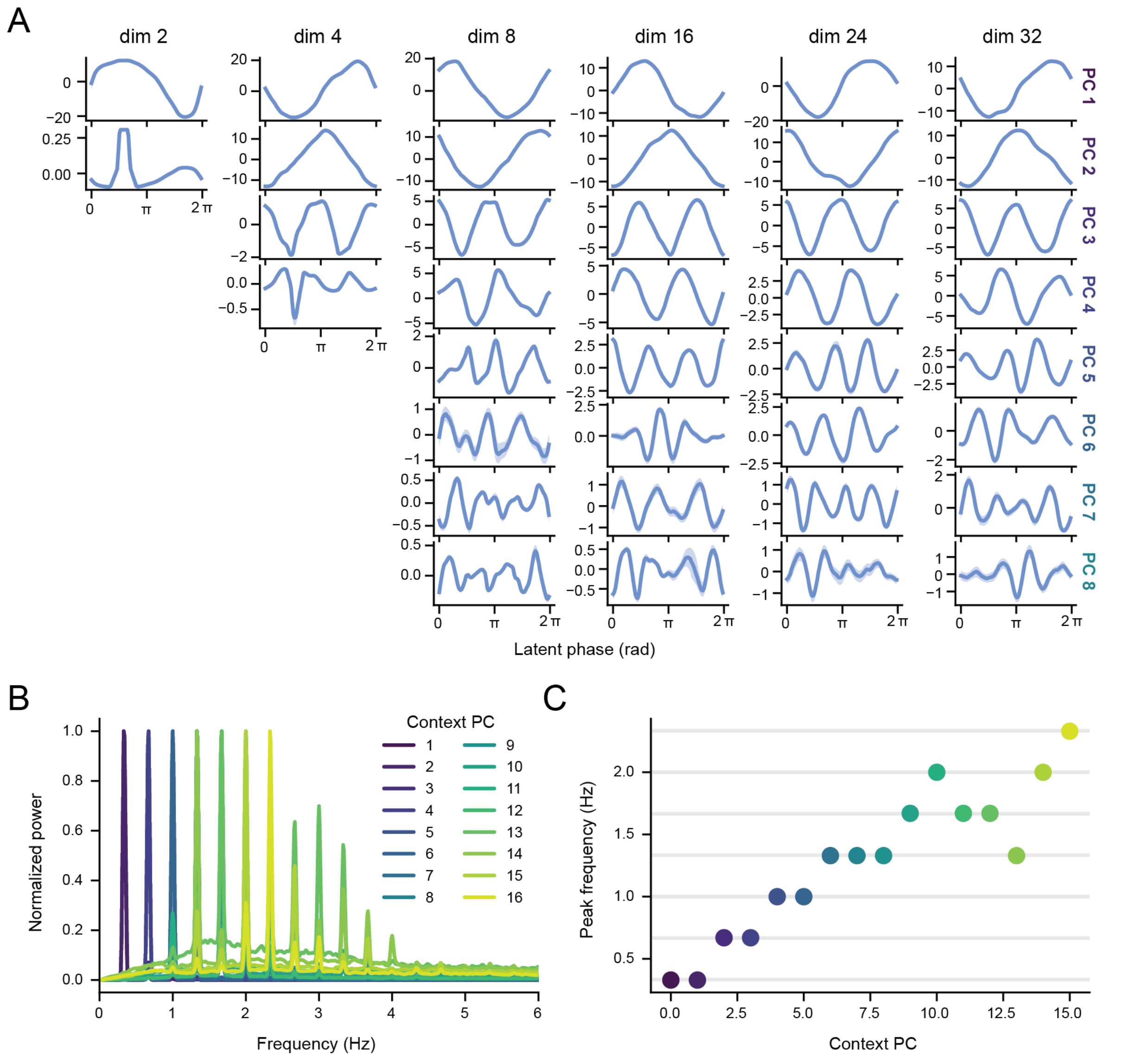


**Supplementary Figure 2.** | **Model dimensionality and harmonic structure of C3PO representations of a periodic latent variable.**

**A,** Context representations from C3PO models with 2, 4, 8, 16, 24 or 32 embedding and context dimensions, independently trained on the same synthetic dataset used in Fig. 1. Each column shows one model; rows show the first eight principal components (PCs), or all components for models with fewer dimensions. Lines and shading indicate median and interquartile context values within latent-phase bins, respectively. PCA was performed separately for each model.

**B,** Frequency-weighted power spectra of all 16 context PCs from the model analyzed in Fig. 1. Spectra were estimated using Welch's method with 30-s Hamming windows and 50% overlap, multiplied by frequency and normalized to their respective maxima. Colors identify individual PCs.

**C,** Frequency of maximal power in each normalized spectrum shown in B. Horizontal gray lines indicate integer multiples of the true latent frequency (one-third hertz, 3 s period). Dominant frequencies align with the fundamental frequency and its higher harmonics, consistent with a Fourier decomposition of the latent phase.

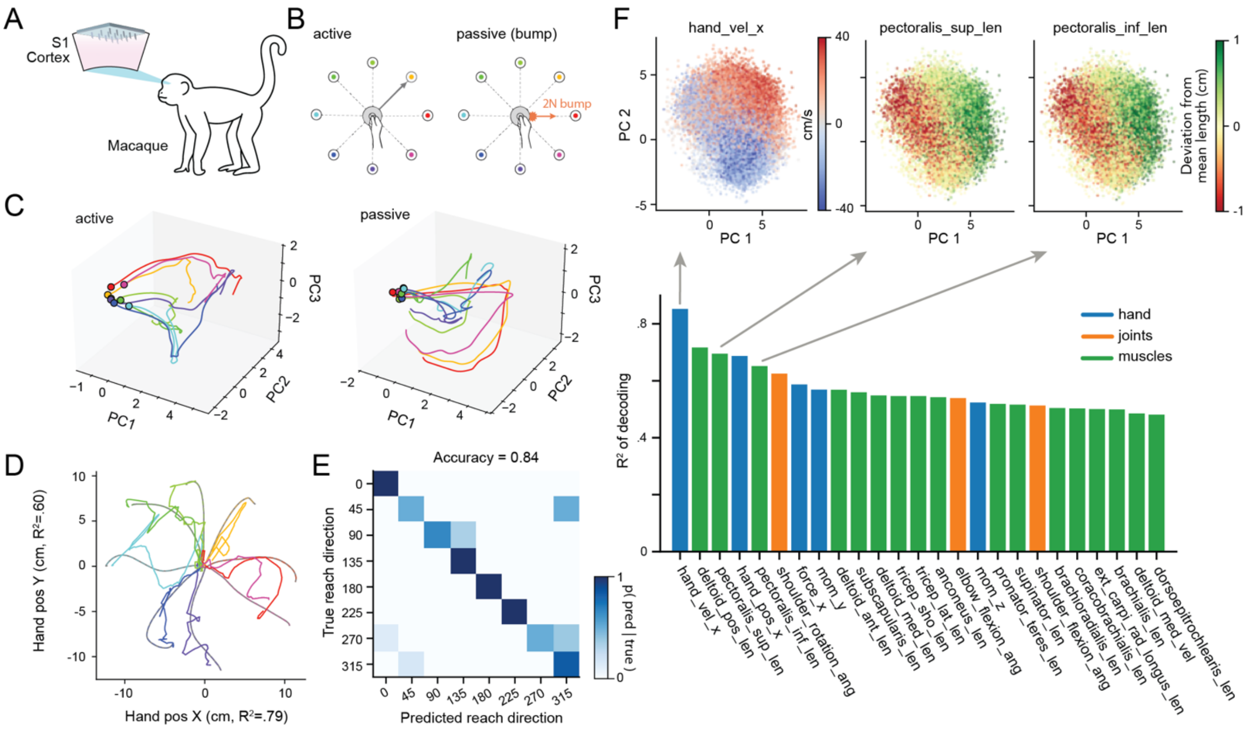


**Supplementary Figure 3.** | **C3PO captures movement and proprioceptive structure in macaque somatosensory cortex.** A 16-dimensional C3PO model was trained on sorted spike events, with neuron identities supplied as one-hot categorical inputs to the mark-embedding network (Methods; Table S1). Behavioral variables and movement labels were used only for subsequent characterization and supervised decoding of the learned context. All decoders were fit and evaluated using the dataset-defined training and validation trial assignments.

**A,** Recording schematic showing a Utah array targeting the arm representation region of primary somatosensory cortex (S1, area 2) in one rhesus macaque[20].

**B,** Active and passive movement conditions. The animal voluntarily reached toward one of eight peripheral targets or experienced an uncued, 2-N mechanical perturbation of the hand. Colors indicate movement direction. Positions of markers along the arm provided the measurements used to derive hand, joint, and muscle-related kinematic variables.

**C,** Mean context trajectories projected onto the first three principal components (PCs), grouped by movement direction for active reaches (left) and passive perturbations (right). Context was aligned to movement onset, averaged over trials from 100 ms before to 500 ms after onset, and Gaussian-smoothed (100 ms standard deviation). Circles mark the beginning of each displayed trajectory; colors follow **B**. Active-reach trajectories originated from a similar pre-movement region before diverging along distinct paths for the eight reach directions. Passive perturbations also produced direction-dependent trajectories that differed from those during voluntary movement. This organization closely replicated movement-related structure previously reported using CEBRA with behavioral supervision or experimenter-defined temporal positive pairs.

**D,** Observed hand trajectories (gray, with dashed colored overlays) and trajectories decoded from C3PO context (solid colored lines) for eight example active validation trials. Predictions were obtained using a distance-weighted nearest-neighbor decoder with Gaussian-smoothed context (100 ms standard deviation). Decoded trajectories closely followed observed reaches across movement directions. Squared Pearson correlations between observed and predicted positions, pooled across validation trials, were 0.79 and 0.60 for horizontal and vertical position, respectively.

**E,** Reach-direction decoding from context averaged from 100 ms before to 500 ms after movement onset, using multinomial logistic regression. The confusion matrix shows prediction proportions normalized within each true direction; overall accuracy was 84% across 51 validation trials, with most errors occurring between nearby directions. Binary logistic regression applied to the same trial-averaged representation distinguished active from passive movements with 94% accuracy on held-out trials (not shown). These decoding accuracies were comparable to those previously reported for supervised CEBRA models.

**F,** Bottom, decoding performance for the 25 highest-scoring hand-related (blue), joint (orange), and muscle-derived (green) variables, quantified by squared Pearson correlation across active validation trials. Nearest-neighbor decoders used context smoothed with a 40-ms Gaussian standard deviation. Top, context projected onto its first two PCs during the first 500 ms of active reaches, colored by horizontal hand velocity or deviations from mean superior and inferior pectoral muscle lengths. Arrows identify the corresponding variables below. The high decodability of these biomechanically related variables and their smooth gradients across latent space indicate that the dominant context axes track coordinated changes in endpoint movement and shoulder configuration. This organization is consistent with the whole-arm proprioceptive representation reported in the original study[20], recovered here without specifying a candidate behavioral encoding model during representation learning.

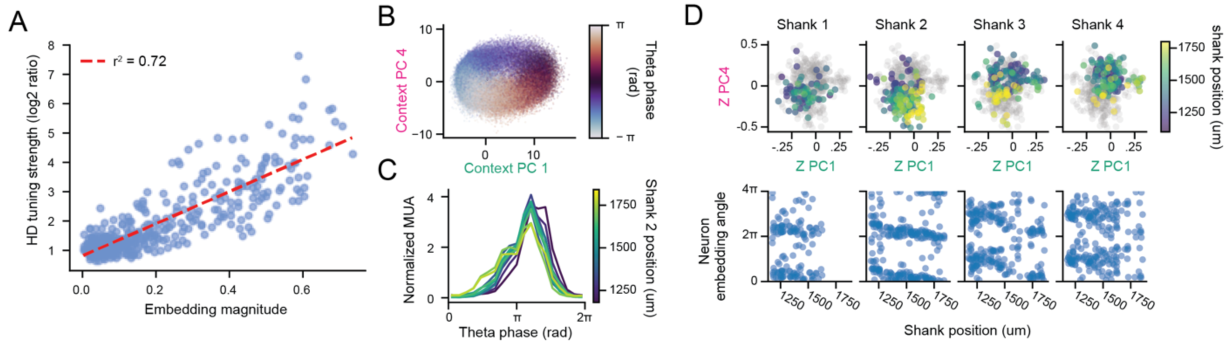


**Supplementary Figure 4. | C3PO embeddings reflect head-direction tuning strength and anatomically organized theta timing.** Analyses use the same recording of 460 neurons from medial entorhinal cortex (MEC) and adjacent parasubiculum in one freely moving rat as Fig. 3.

**A,** Relationship between neuronal embedding magnitude in the head-direction-sensitive PC2–PC3 plane and observed head-direction tuning strength. Tuning strength is the base-2 logarithm of the ratio between peak and median firing rates across head-direction bins. Each point represents one neuron. The red dashed line shows a least-squares linear fit ($R^2 = .71$). Neurons with stronger head-direction modulation have larger embedding magnitudes, consistent with the model's mapping between embedding weights and firing-rate modulation.

**B,** C3PO context projected onto the theta-sensitive PC1–PC4 plane, colored by the independently measured theta phase. Theta phase varies continuously around the context distribution, revealing a circular organization of the population representation.

**C,** Theta-phase dependence of normalized multiunit activity (MUA) at different recording positions along shank 2. Neurons were grouped into ten equal-width position bins, and their firing rates were summed within each group. Colors indicate position along the shank. Theta phase is offset by half a cycle for display. Position-dependent differences in the timing and shape of these profiles indicate depth-dependent offset in theta phase.

**D,** Anatomical organization of neuronal embeddings in the theta-sensitive plane. Top, embeddings projected onto the context PC1–PC4 axes, shown separately for each probe shank. Colored points represent neurons on the indicated shank, colored by recording position; gray points show the full population for reference. Bottom, embedding angle in this plane plotted against position along the corresponding shank. Angles are repeated over two cycles to display circular continuity. The relationship between embedding angle and recording position reveals anatomical structure in the learned theta-related neuronal weights.

Together, **B–D** support the interpretation that C3PO captures anatomically organized differences in relative neuronal firing across theta phase. Importantly, because C3PO aims to identify features that differentially predict which specific spike waveform or neuron will fire next, modulations common to all units will not be directly detected. In this case C3PO identifies theta because different neurons are most active at different theta phases, and thus theta phase is highly informative as to which neurons will be preferentially active at a given time. This interpretation is consistent with previously reported laminar organization of MEC theta fields and neuronal phase preferences (Mitchell and Ranck, 1980; Mizuseki et al., 2009; Quilichini et al., 2010), and with progressive theta-phase shifts in field activity and multiunit firing along the dorsoventral axis (Hernández-Pérez et al., 2020).

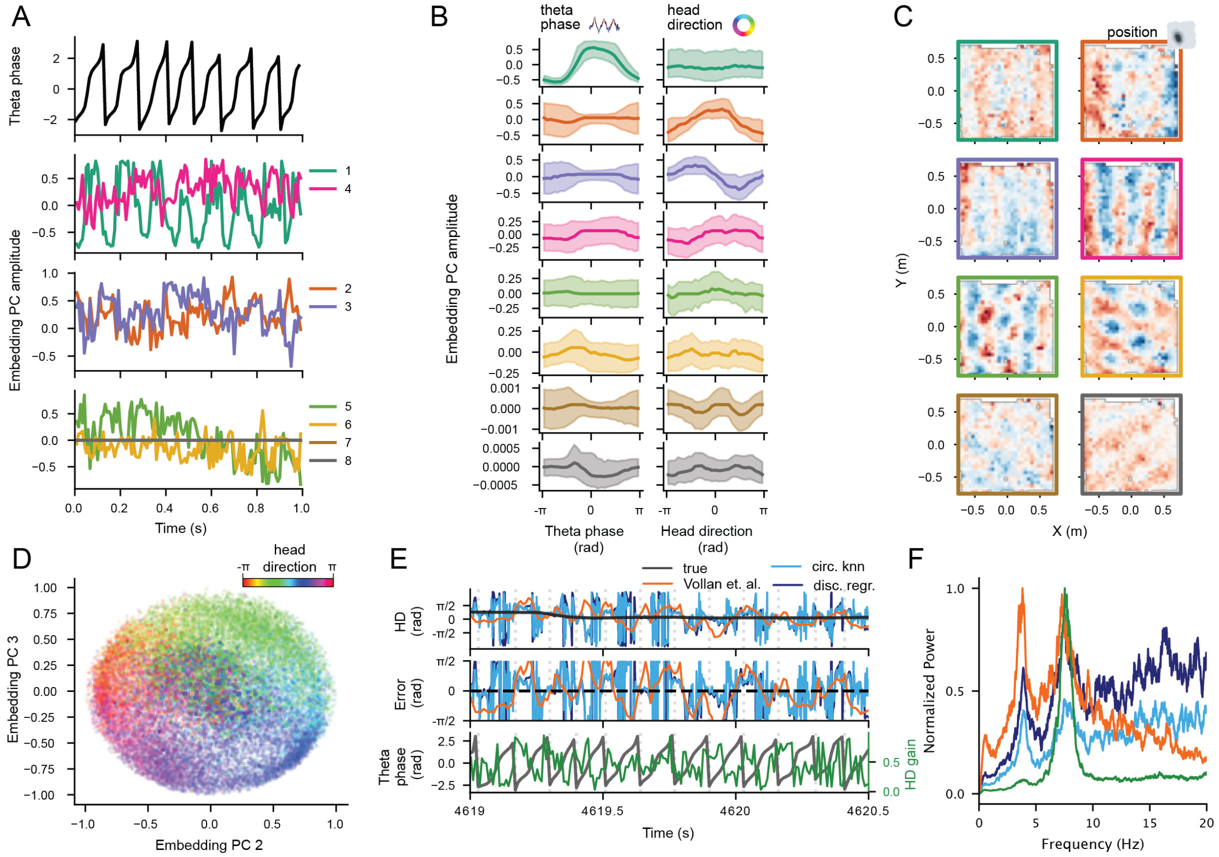


**Supplementary Figure 5. | CEBRA embeddings of entorhinal population activity.** CEBRA was applied to the same Neuropixels recording of 460 sorted neurons from rat medial entorhinal cortex (MEC) and adjacent parasubiculum analyzed in Fig. 3 (Vollan et al., 2025). Analyses follow those in Fig. 3, using principal components (PCs) of the CEBRA latents in place of C3PO context.

**A,** Example trace of theta phase (top) and the first eight PCs of the CEBRA latents (below). Colors identify PCs throughout **A–C**.

**B,** Latent-PC values conditioned on theta phase (left) or head direction (right). Lines and shading indicate median and interquartile values, respectively.

**C,** Spatial maps of median latent-PC values for PCs 1–8; border colors match **A,B**. Each map uses a symmetric color scale spanning its maximum absolute value, with red and blue indicating negative and positive values, respectively.

**D,** CEBRA latents projected onto PCs 2 and 3, colored by measured head direction. Head-direction gain denotes the radial magnitude within this plane.

**E,** Example head-direction dynamics. Top, measured head direction (gray), five-fold cross-validated CEBRA predictions from multinomial logistic regression (dark blue), circular nearest-neighbor regression (light blue), and the internal-direction estimate supplied by Vollan et al. (orange). Middle, signed circular decoding errors relative to measured head direction. Bottom, theta phase (gray) and head-direction gain (green). Vertical dashed lines mark theta-phase cycles.

**F,** Frequency-weighted power spectra of signed decoding error and head-direction gain (green), each normalized to its maximum. Decoder colors follow **E**.

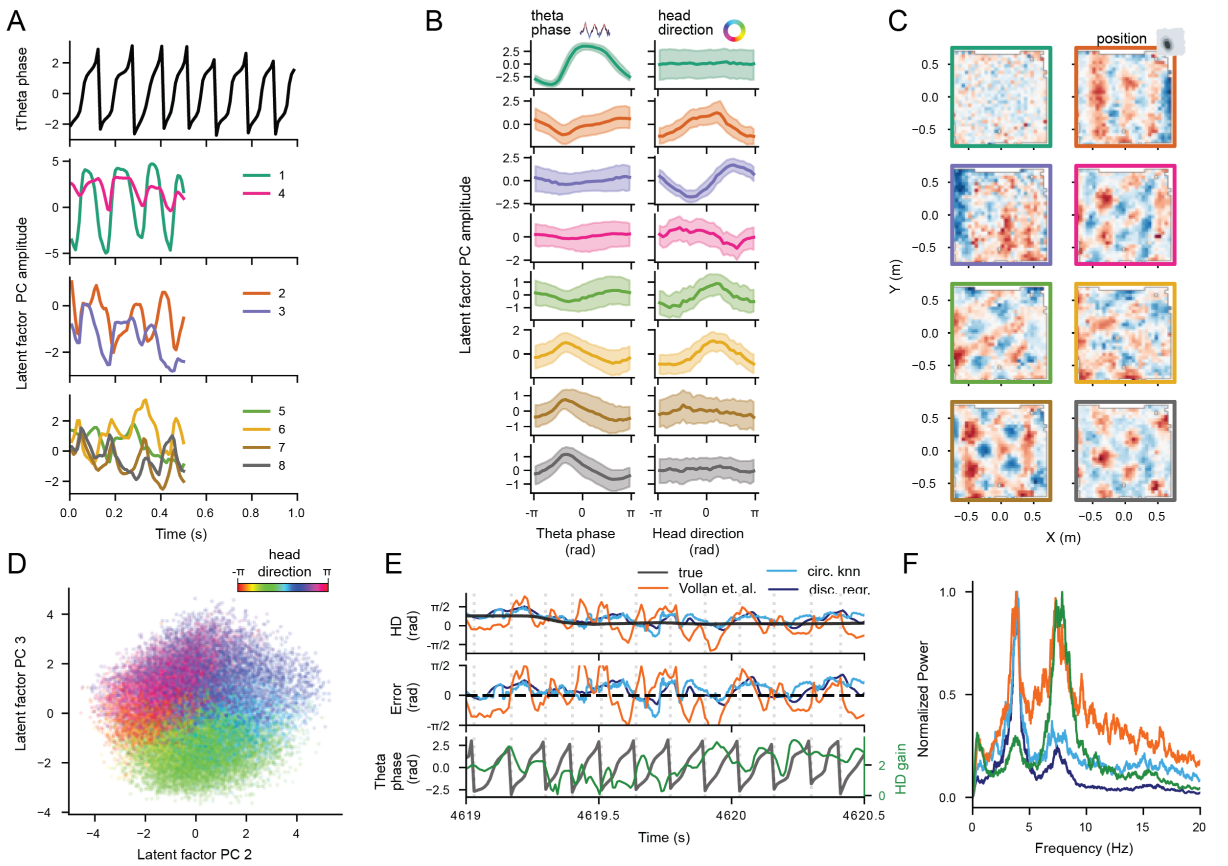


**Supplementary Figure 6.** | **LFADS latent factors from entorhinal population activity.** LFADS was applied to the same Neuropixels recording of 460 sorted neurons from rat medial entorhinal cortex (MEC) and adjacent parasubiculum analyzed in Fig. 3 (Vollan et al., 2025). Analyses follow those in Fig. 3, using principal components (PCs) of the LFADS latent factors in place of C3PO context.

**A,** Example trace of theta phase (top) and the first eight PCs of the LFADS latent factors (below). Colors identify PCs throughout **A–C**. LFADS factor trajectories are truncated because of padding introduced to accommodate gaps in the public dataset.

**B,** Factor-PC values conditioned on theta phase (left) or head direction (right). Lines and shading indicate median and interquartile values, respectively.

**C,** Spatial maps of median factor-PC values for PCs 1–8, arranged in row-major order; border colors match **A,B**. Each map uses a symmetric color scale spanning its maximum absolute value, with red and blue indicating negative and positive values, respectively.

**D,** LFADS factors projected onto PCs 2 and 3, colored by measured head direction. Head-direction gain denotes the radial magnitude within this plane.

**E,** Example head-direction dynamics. Top, measured head direction (gray), five-fold cross-validated LFADS predictions from multinomial logistic regression (dark blue), circular nearest-neighbor regression (light blue), and the internal-direction estimate supplied by Vollan et al. (orange). Middle, signed circular decoding errors relative to measured head direction. Bottom, theta phase (gray) and head-direction gain (green). Vertical dashed lines mark theta-phase cycles.

**F,** Frequency-weighted power spectra of signed decoding error and head-direction gain (green), each normalized to its maximum. Decoder colors follow **E**.

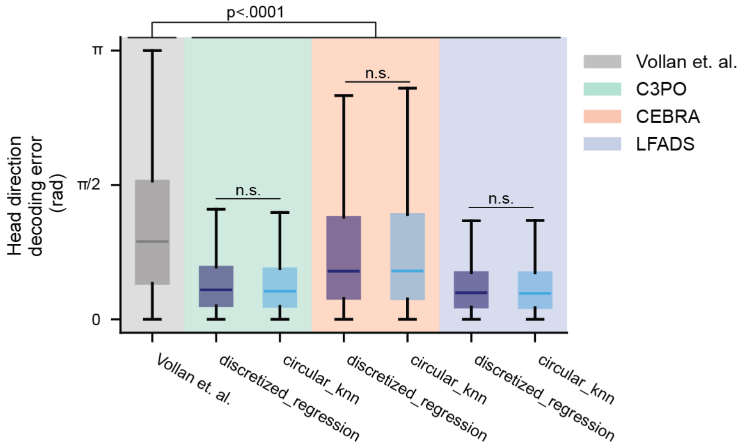


**Supplementary Figure 7.** | **Head-direction decoding performance across embedding methods.** Distributions of absolute circular decoding error for the internal-direction estimate supplied by Vollan et al. (gray) and predictions from C3PO context, CEBRA latents and LFADS latent factors. Data are from the rat MEC and adjacent parasubiculum recording analyzed in Fig. 3. For each embedding method, multinomial logistic regression (dark blue) and circular nearest-neighbor regression (light blue) were evaluated using five-fold temporal cross-validation. Error is the magnitude of the shortest angular difference between predicted and measured head direction. Background shading identifies the embedding method. Boxes indicate interquartile ranges, center lines indicate medians and whiskers extend to the most extreme values within 1.5 interquartile ranges of the box.

Pairwise differences in mean error were evaluated using two-sided permutation tests that shuffled mean errors from nonoverlapping 0.5-s windows within continuous recording intervals between comparison groups (10,000 permutations per comparison). Each of the six decoders yielded lower mean error than the Vollan et al. estimate (top bracket, all $p<.0001$). For both decoder types, mean error was lowest for LFADS, followed by C3PO and CEBRA (all pairwise comparisons between embedding methods for a given decoder type, all $p<.0001$). Within each embedding method, the difference between decoder types was nonsignificant (n.s., $p = .1034$, *.0952, and* .6987 for C3PO, CEBRA and LFADS, respectively).

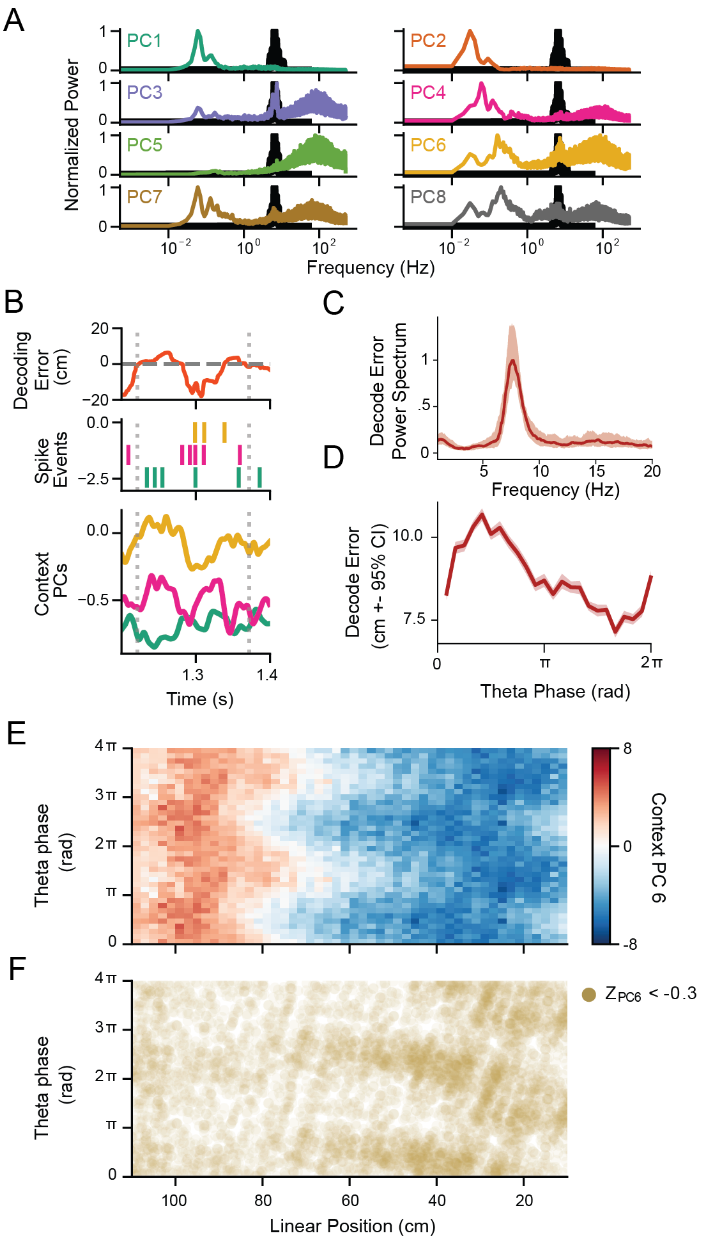


**Supplementary Figure 8.** | **Theta-timescale sequences and phase precession in clusterless hippocampal representations.** Data are from the CA1 recording shown in Fig. 4.

**A,** Power spectra of the first eight context PCs with the theta-band local field potential (LFP) spectrum overlaid in black. Spectra were estimated using Welch's method, multiplied by frequency and independently normalized to their maxima.

**B,** Expanded view of the traversal shown in Fig. 4D–H, illustrating sequential context modulation within a theta cycle. Top, signed C3PO position-decoding error. Middle, clusterless events with strongly negative waveform-embedding projections onto context PC6 (gold), PC4 (magenta), or PC1 (green), selected using the lowest 2.5% of projections along each axis during the complete trial. Bottom, corresponding context components, smoothed and normalized to their maximum absolute values during the traversal. Vertical dotted lines mark theta-LFP maxima.

**C,** Power spectrum of signed position-decoding error during running. The curve shows the median and interquartile range of the spectrum across running intervals, multiplied by frequency and normalized to its peak. The peak near 8 Hz reflects theta-frequency fluctuations in decoded position relative to measured position.

**D,** Mean absolute position-decoding error as a function of theta phase during running. Shading indicates 95% bootstrap confidence intervals calculated from 1,000 resamples of timepoints within each phase bin.

**E,** Mean context PC6 as a function of linear position and theta phase during leftward running. Diagonal structure indicates changing phase dependence of the context across position.

**F,** Theta phase and position of individual clusterless events whose waveform embeddings project below −0.3 onto context PC6. The position-dependent phase distribution parallels the context structure in E and is consistent with phase precession in an event group identified without spike sorting. In E and F, position decreases from left to right, following the direction of travel, and the same data are repeated over two theta cycles to display circular continuity.

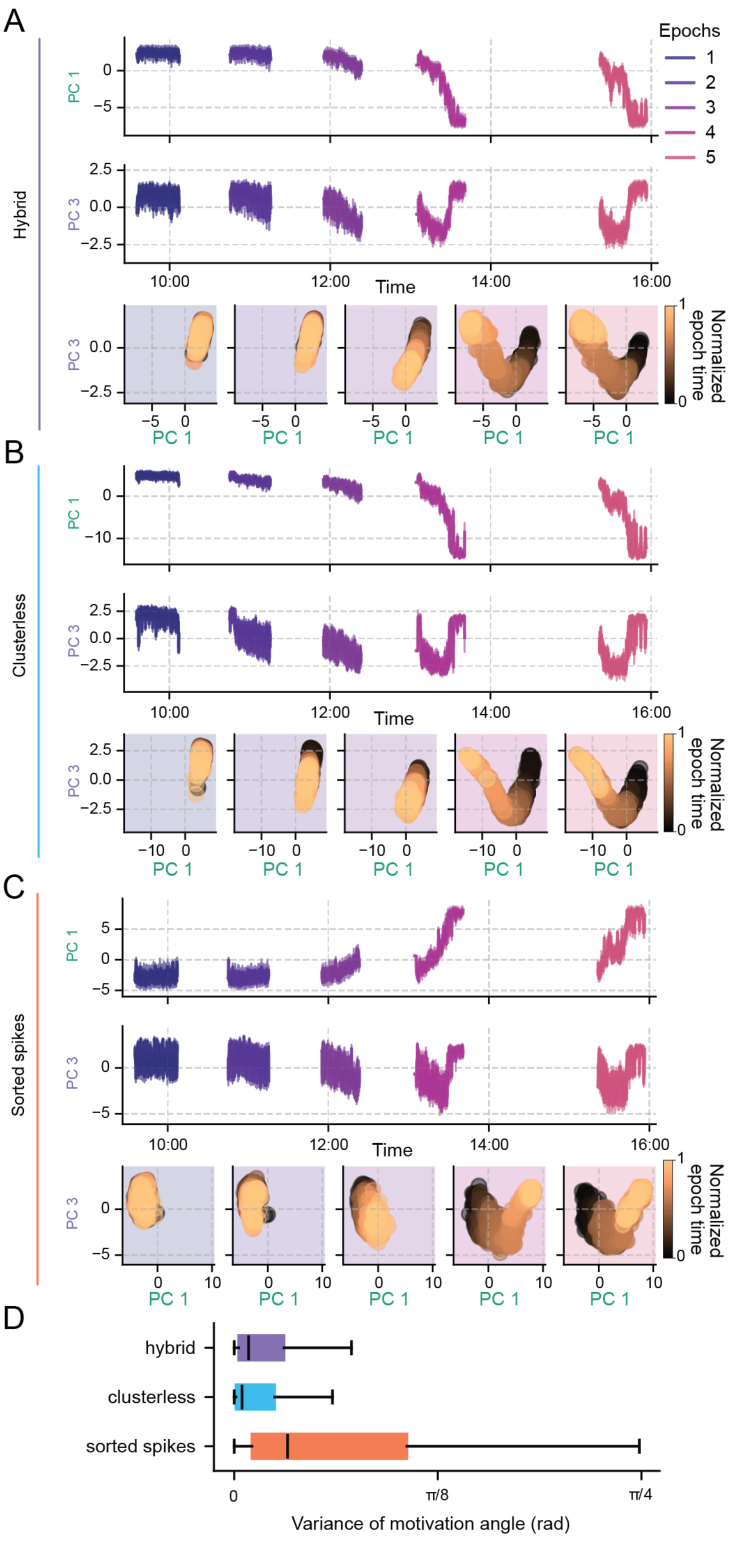


**Supplementary Figure 9.** | **Slow prefrontal state dynamics are preserved across spike representations.** C3PO models were trained independently using three representations of the same mPFC.

**A,** Hybrid model combining neuron-specific embeddings of sorted spikes with waveform-based embeddings of unassigned events, as in Fig. 5. Curves show medians within 3-s windows; shading indicates interquartile ranges. Gaps separate task epochs interleaved with rest. The lower plots show the corresponding context trajectories in the PC1–PC3 plane, separately for epochs 1–5 from left to right. Points represent window medians and are colored by normalized time within each epoch, from black at the beginning to light orange at the end. Principal components were estimated separately for each model, allowing differences in axis sign and scale.

**B,** Model trained using clusterless waveform features and event times.

**C,** Model trained using sorted spike times and neuron identities alone. All three models recover prominent slow excursions late in epochs 4 and 5 and a return toward earlier states after intervening rest. The shared temporal organization supports the robustness of the engagement-associated population state, operationally termed "motivation," across hybrid, sorted-only, and clusterless input representations.

**D,** Distribution of the variance of the estimated motivation angle within each 3s window shown in **A-C**. Models including clusterless marks in the input data showed significantly lower variance in the motivation angle estimate than the model trained on sorted spike data alone (permutation of mean values, $p < 1e - 5$ for both hybrid and clusterless model). This suggests that the inclusion of spike amplitudes and low-confidence events allows C3PO to extract more informative latent variables corresponding to the internal state.
